# Crop modeling defines opportunities and challenges for drought escape, water capture, and yield increase using chilling-tolerant sorghum

**DOI:** 10.1101/2021.01.27.428532

**Authors:** Rubí Raymundo, Sarah Sexton-Bowser, Ignacio A. Ciampitti, Geoffrey Morris

## Abstract

Many crop species, particularly those of tropical origin, are chilling sensitive so improved chilling tolerance can enhance production of these crops in temperate regions. For the cereal crop sorghum (*Sorghum bicolor L.*) early planting and chilling tolerance have been investigated for >50 years, but the potential value or tradeoffs of this genotype × management change has not been formally evaluated with modeling. To assess the potential of early-planted chilling-tolerant grain sorghum in the central US sorghum belt, we conducted CERES-Sorghum simulations and characterized scenarios under which this change would be expected to enhance (or diminish) drought escape, water capture, and yield. We conducted crop growth modeling for full- and short-season hybrids under rainfed systems that were simulated to be planted in very early (April), early (May 15), and normal (June 15) planting dates over 1986–2015 in four locations in Kansas representative of the central US sorghum belt. Simulations indicated that very early planting will generally lead to lower initial soil moisture, longer growing periods, and higher evapotranspiration. Very early planting is expected to extend the growing period by 20% for short- or full-season hybrids, reduce evaporation during fallow periods, and increase plant transpiration in the two-thirds of years with the highest precipitation (mean > 428 mm), leading to 11% and 7% increase grain yield for short- and full-season hybrids, respectively. Thus, in this major sorghum growing region very early and early planting could reduce risks of terminal droughts, extend seasons, and increase rotation options, suggesting that further development of chilling tolerant hybrids is warranted.

## INTRODUCTION

Sustainable agriculture in water-limited environments requires cropping systems with low water requirements and relatively high productivity per unit of water used (Tilman et al., 2002; Zhang et al., 2018). Breeding programs generally rely on multi-environment trials in the target environment to assess the performance of genotypes and evaluate novel traits. However, this approach has limitations: (i) resources to deploy such trials are often limited, (ii) it is difficult to reproduce and/or interpret findings from year to year due to spatio-temporal variability of environmental conditions, and (iii) in most of the cases the novel trait is not yet available in commercial varieties, so cannot be studied in relevant genetic backgrounds (Lenaerts et al., 2019). In this context, crop growth models complement field experiments (van Ittersum et al., 2003; Challinor et al., 2018) to help breeding programs evaluate the potential value of novel traits within an existing or novel cropping system (Aggarwal et al., 1997; Chenu et al., 2017; Cooper et al., 2002). Models can be used to evaluate the potential benefits and tradeoffs of combining new genotypes and new management approaches in the target production environments, by accounting for interactions of genotype, environment and management (G×E×M) (Chenu et al., 2013; Kholová et al., 2014; Singh et al., 2014).

Sorghum (*Sorghum bicolor* L. Moench) is one of the most drought resilient crops, grown worldwide for grain, forage, and biomass (Adegbeye et al., 2020). The crop originated in the warm semi-arid tropics of Africa (Doggett and Majisu, 1968) and diffused widely, including to semi-arid and sub-humid temperate regions of the world (Smith and Frederiksen, 2000). However, like many tropical-origin crops, sorghum is sensitive to chilling (0-16 °C) (Lyons, 1973; Taylor and Rowley, 1971). Chilling sensitivity may be an ancestral trait stemming from sorghum’s origin and adaptation in warm climates, or a derived trait originating inadvertently during selection for non tannin and/or semi-dwarf traits (Marla et al., 2019). Following Vavilov’s phytogeographic approach, seedling chilling tolerance was identified in high-latitude Asian (Stickler et al., 1962) and high-altitude African (Singh, 1985) landraces, and more recently the trait was genetically mapped (Burow et al., 2011; Chopra et al., 2017; Marla et al., 2019). However, this trait has remained a breeding target for >50 years and has not been deployed in commercial grain sorghum hybrids. Research and development on chilling-tolerant sorghum remains ongoing in Europe (Schaffasz et al., 2019), Asia (Mocoeur et al., 2015), Australia (Wylie, 2008), and the Americas (Marla et al., 2019).

In temperate production regions, chilling temperatures restrict the sorghum growing season from late spring to fall (Ercoli et al., 2004). In the semi-arid region of the US sorghum belt (Laingen, 2015), a commercial hybrid with early-chilling tolerance trait has the potential to change the agricultural landscape by shifting planting dates from late spring to early spring to hypothetically position the crop in better environments to ensure early establishment (Burow et al., 2011; Franks et al., 2006; Kapanigowda et al., 2013). In the US, the state of Kansas leads grain sorghum production with 40% of national production on 2.7 million acres, mostly under rainfed conditions (>90%) (Laingen, 2015). Early crop establishment could potentially extend the crop growing period, synchronizing crop water demand and soil water supply during the grain filling to avoid terminal droughts. Furthermore, early planting practices can facilitate integrating the crop into a double crop rotation to maximize farm productivity (Burow et al., 2011). For these reasons, it is believed that the early-season chilling tolerance trait would make the crop more competitive for farmers and an alternative option to outperform traditional crops with high water demand and less adapted to dryland systems (Bhattarai et al., 2019).

Ongoing efforts to improve chilling tolerance in sorghum (Burke et al., 2019; Chopra et al., 2017; Marla et al., 2019; Ostmeyer et al., 2020) are based on the hypothesis that early planting of a sorghum hybrid with chilling tolerance provides a commercially relevant increase in grain yield in the US by extending the season and avoiding late season drought. Due to the lack of chilling tolerant commercial hybrids this hypothesis cannot be directly tested under field conditions. However, this hypothesis can be formally evaluated with crop modeling. In this study we used the CERES-Sorghum growth model (Jones et al., 2003; White et al., 2015) to test these hypotheses *in silico*, and quantify the expected agronomic impact of an early-chilling tolerant sorghum hybrids in the northern part of the U.S. sorghum belt. We find support for several of the proposed benefits of early-planted chilling-tolerant sorghum, along with some unanticipated tradeoffs that must be accounted for in the overall G×E×M strategy for this cropping system.

## MATERIALS AND METHODS

### The sorghum cropping system

Kansas has a precipitation gradient decreasing from east to west (Lin et al., 2017) (Figure 1a), with Koppen-Geiger climate classification of continental-humid in the east and semi-arid climate in the west (Pražnikar, 2017). This gradient in environmental conditions has substantially influenced the management of this rainfed crop such as plant density and varietal selection. Recommended plant density declines from east to west, and later maturity hybrids are recommended in the east while earlier maturity hybrids are recommended in the west (Roozeboom and Fjell, 1998). Given sorghum’s chilling sensitivity and the common practice of prioritizing planting of other crops (i.e maize), sorghum is one of the later-planted crops in the system and consequently the crop can not take advantage of the cumulative growing degrees units or thermal time accumulated in early spring (Figure 1b). The sorghum growing season ranges from early-May to early-July and the main planting season concentrates in the second week of June (Figure 1c) (Shroyer et al., 1998). Otherwise, maize is planted from early-April to mid-June and the main planting season concentrates in mid-May (Figure 1c). Therefore, introgressing chilling tolerance in sorghum would allow the crop to be planted in early spring.

**Figure 1.**
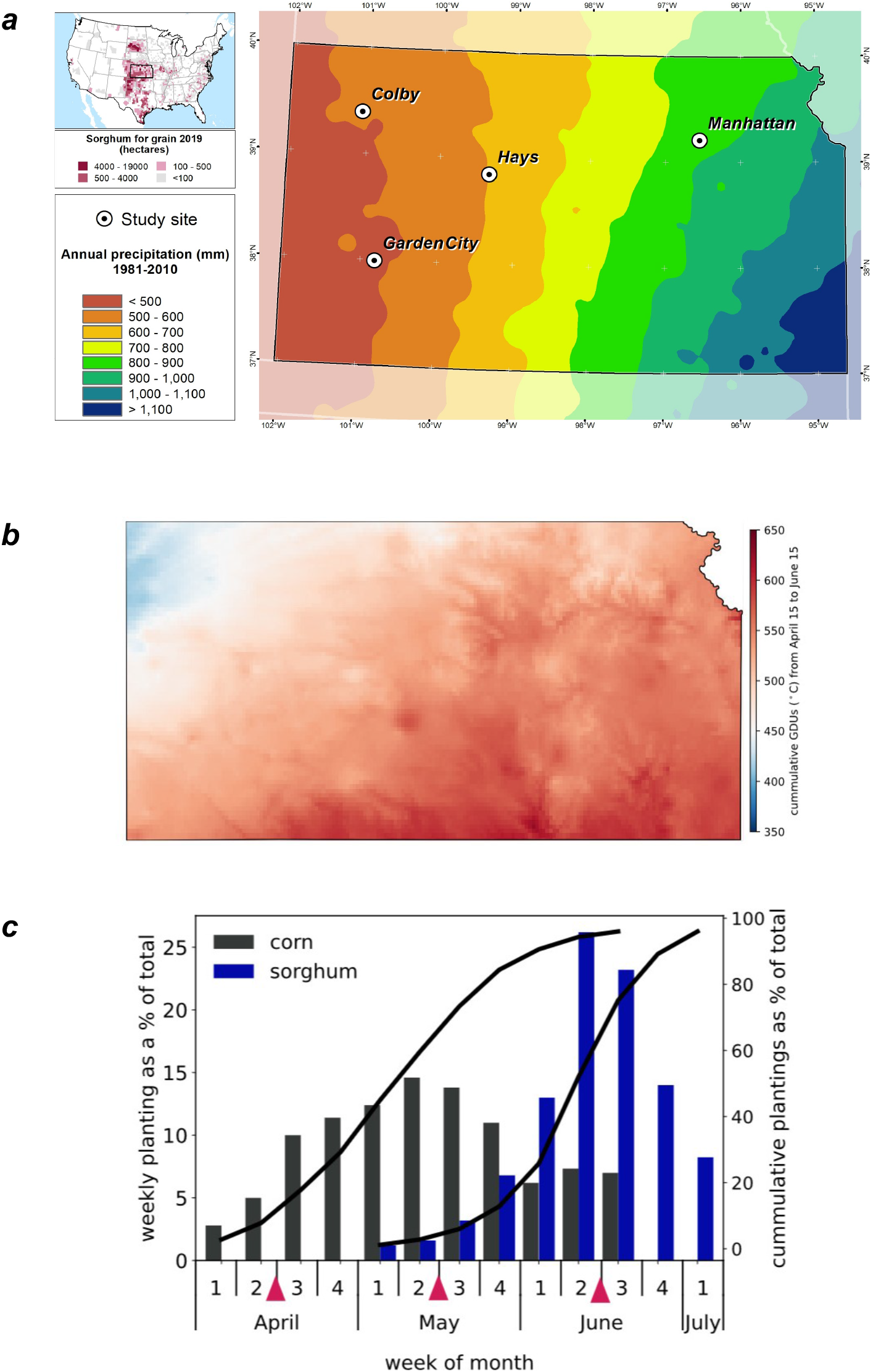
Study system to evaluate the impact of early planting with CERES-Sorghum. (a) Annual precipitation gradients across Kansas and locations to simulate the growth and development for sorghum in Kansas. (b) Cumulative growing degree units (GDU) or thermal time between mid-April to mid-June. The map was generated using the daily average mean over five years of daily data (https://prism.oregonstate.edu/). (c) The rate of sorghum and plantings in Kansas, barplots represent the five-year mean (2010-2016), black lines represent the cumulative planting (https://www.nass.usda.gov/) and the pink triangles represent the planting dates under study: very early (April 15), early (May 15), and normal (June 15).

### Overview of the model

CERES-Sorghum (White et al., 2015) belongs to the family of crop models available in the Decision Support Systems for Agro technology Transfer-Crop Simulation Model (DSSAT-CSM) (Hoogenboom et al., 2019; Jones et al., 2003). The model simulates the daily dynamics of plant growth, phenology development and partitioning affected by stress factors (White et al., 2015).

The daily growth is simulated via the potential carbon assimilation (PCARB) that is calculated using the radiation use efficiency (RUE, g MJ^−1^), photosynthetically active radiation (PAR, MJ m^−1^ day^−1^) and the fraction of PAR intercepted (1 - exp^(−k * LAI)^) by the crop (Ritchie et al., 1998), where k (unitless) represents the extinction coefficient and LAI (m_leaf_^2^ m_ground_^−2^) is the leaf area index. The model simulates daily water stress factors that penalize the daily carbon assimilation (SWFAC) and plant growth (TURFAC) (White et al., 2015). The model requires cultivar-specific genotype parameters (G), environmental information (E), and crop management (M) practices, which are described in detail below.

### Genotype parameters

In the model, G, a set of genotype-specific parameters, represents differences among hybrids (White et al., 2015) that affect phenology development and crop growth. In this study, the difference between early and late season hybrids is represented by the parameter potential duration from emergence to end of juvenile phase (P1) (Table 1). In the model the phenology development is a function of temperature and daylength. The model simulates germination, emergence, end of the juvenile phase, panicle initiation, end of flag leaf expansion, anthesis, beginning of grain filling and physiological maturity. Each phenology stage is the result of the summation of daily thermal time (DTT) calculated via the formula

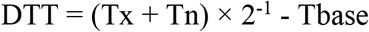

where Tbase is the minimum temperature for thermal time accumulation, set to 8 °C (White et al., 2015), and Tx and Tn are daily maximum and minimum temperature, respectively. Note, given that the DTT uses a Tbase of 8 °C and does not implement a chilling damage routine (i.e. there is no penalty to growth or development for temperatures below Tbase) we interpret CERES-Sorghum as modeling a chilling-tolerant sorghum genotype by default. The summation of DTT to reach a phenology stage is indicated in Table 1 (e.g. P1: Potential duration from emergence to end of juvenile phase). The only phenology stage affected by both temperature and daylength is the duration from the end of juvenile stage to panicle initiation, represented by parameters P2R and P2O (Table 1). The plant leaf growth and panicle partitioning are determined by parameter G1 and G2, respectively.

**Table 1.** Cultivar parameters for CERES-Sorghum model for full- and short-season hybrids*.

| **Code** | **Full**<br>**season** | **Short**<br>**season** | **Description** |
| --- | --- | --- | --- |
| P1 | 393 | 250 | Potential duration from emergence to end of juvenile phase (°C day). |
| P2 | 102 | 102 | Potential duration from juvenile phase to panicle initiation (°C day). |
| P2O | 15.0 | 15.0 | Critical daylength above which development slows (short day response) (h). |
| P2R | 40 | 40 | Photoperiod sensitivity as the extent to which development is delayed for each hour of photoperiod above P2O (°C day). |
| PANTH | 617 | 617 | Duration from panicle initiation to anthesis (°C day). |
| P3 | 152 | 152 | Duration from end of flag leaf expansion to anthesis (°C day). |
| P4 | 82 | 82 | Duration from anthesis to onset of grain filling (°C day). |
| P5 | 640 | 640 | Duration of grain-filling phase (onset of grain filling to physiological maturity) (°C day). |
| PHINT | 49 | 49 | Phyllochron interval (°C day/leaf). |
| G1 | 3 | 3 | Scaler for relative leaf size (unitless). |
| G2 | 6.5 | 6.5 | Scalar for partitioning of assimilates to the panicle (unitless). |

\* Reproduced from Araya et al. (2018).

**Table 2.** Characteristics for the study locations in Kansas.

| **Location** | **Soil family** | **Soil texture** | **Organic matter (%)** | **Average annual precipitation (mm)** | **Koppen Climate classification** |
| --- | --- | --- | --- | --- | --- |
| Colby | Richfield | Silty clay loam | 2 | 520 | Cold semi-arid |
| Garden City | Park-Ulysses | Silt loam | 2 | 480 | Cold semi-arid |
| Hays | Harney | Silt loam | 2 | 680 | Hot-summer humid continental |
| Manhattan | Smolan | Silt loam | 3 | 900 | Hot-summer humid continental |

### Environmental data

In the model, E comprises daily weather data and multilayer soil profile parameters. Daily weather data, including precipitation (mm), solar radiation (MJ m^−2^ day^−1^), and daily maximum (°C) and minimum temperature (°C) was obtained for agronomic research sites (Manhattan, Hays, Garden City, and Colby) from Kansas Mesonet from 1986 to 2015 (Kansas Mesonet, 2020). Soil profile information was downloaded from the web soil survey (Soil Survey Staff, 2020) and detailed soil characteristics such as soil texture (%), bulk density (g ml^−1^), organic carbon (%), pH, wilting point (LL), field capacity (DUL), saturation (SAT), and soil root growth factor (SRGF) for each site are presented in Table S1. The model computes the differences between DUL and LL, and between SAT and DUL, to predict the daily soil extractable water and the water drained in each soil layer, respectively. And, the SRGF represents the physical or chemical constraints in each soil profile.

### Management practices

The model requires realistic agronomic management (M) data such as planting date, planting density, row spacing, fertilization, and water supply. In this study, the crop was simulated to be planted each year in April 15 (very early), May 15 (early), and June 15 (normal) at a row distance of 76 cm, a planting depth of 3 cm, and nitrogen fertilizer rates of 190 kg ha^−1^. Plant density was 14 plants m^−2^ in Manhattan, 8 plants m^−2^ in Hays, and 6 plants m^−2^ in both Garden City and Colby. Given that sorghum grows mostly under dryland systems across Kansas, simulations were performed under rainfed conditions.

### Seasonal simulation settings

As outlined above, information on G, E, and M were defined for each research site. Simulations under rainfed conditions started on the first day of each year with 80% of soil moisture, 3 Mg ha^−^ 1 of surface residue, and 2 Mg ha^−1^ of root residue. Evapotranspiration was calculated with the Priestley-Taylor/Ritchie formula (Priestley and Taylor, 1972). Soil water infiltration was computed with the capacity approach method (Ritchie et al., 1998) and soil evaporation was estimated with the Suleiman-Ritchie method (Suleiman and Ritchie, 2003). Dynamics of carbon and nitrogen were simulated with the CENTURY model.

### Evaluation of model performance and the effect of Tbase on days to emergence

For normal planting dates (June), as shown in Figure 2, CERES-Sorghum has shown satisfactory predictions for anthesis, grain yield, and evapotranspiration across Kansas (Araya et al., 2017; Staggenborg and Vanderlip, 2005; White et al., 2015) using a Tbase of 8 °C. To demonstrate that simulations in early spring represent a chilling-tolerant sorghum, the model accuracy in predicting days to emergence for a chilling-tolerant genotype (Kaoliang), simulations were conducted to match field experiments using a Tbase of 6 °C, 8 °C, and 10 °C. In these scenarios, we expect that seedling emergence for simulations with a Tbase of 8 °C will be closer to field data. Simulations were set up for field experiments that were planted in April, May, and June in 2016–2018 at three sites in Kansas (Marla et al., 2019). For these simulations, cultivar specific-parameters (G) were obtained for full-season hybrids (80 days to anthesis and 140 days to physiological maturity) (Araya et al., 2018). The model accuracy was quantified using the root mean square error (RMSE) that determines the distance of model prediction from a perfect prediction (Wallach et al., 2014).

**Figure 2.**
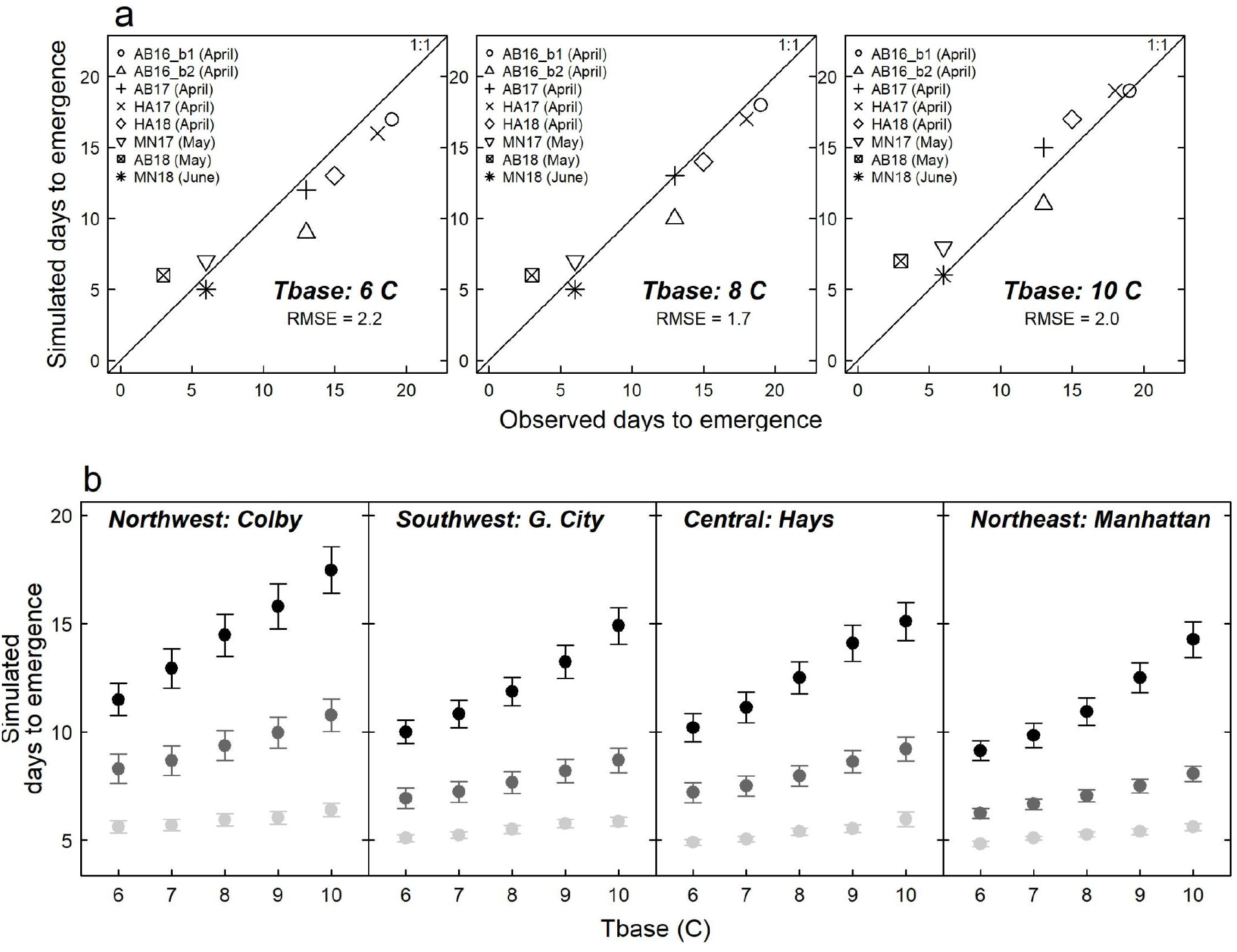
CERES-Sorghum model performance for days to seedling emergence. (a) CERES-Sorghum model performance for days to seedling emergence in different planting dates in Kansas for Tbase of 6 °C, 8 °C, and 10 °C, observed information corresponds to experiments conducted in Hays (HA), Manhattan (MA), and Ashland Bottom (AB; 10 km from MA) in 2016–2018 that were planted in April, May and June. (b) Variation in simulated days to emergence as a function of Tbase in very early (April 15; black points), early (May 15, dark gray points), or normal (June 15, light gray points) planting. Each point represents the mean over 30 years and error lines indicate the standard error.

Unlike commercial grain sorghum hybrids with semi-dwarf stature and high sink capacity, kaoliang sorghum has undesirable traits for this cropping system such as tall plants and low harvest index (Marla et al., 2019). Due to these differences, the direct comparison of observed and simulated grain weight for these known chilling tolerant genotypes is not possible. To ensure that a Tbase of 8 °C represents a sorghum tolerant to chilling temperatures, when planted early, a sensitivity analysis was conducted for a Tbase ranging from 6 °C to 10 °C. Under this assumption, in the simulations the crop will require less time for seedling emergence in early planting dates. For sensitive analysis, simulations were conducted under well-watered conditions to remove the confounding effect of soil moisture on emergence. In our simulations the genotype used across all planting dates have the same cultivar parameters (Table 1) so the effect of planting date can be isolated.

### Statistical analysis and interpretation

Statistical analyses were performed in the R statistical environment (R Core Team, 2017) with lmer (Bates et al., 2015) for mixed linear models and visualized using ggplot (Villanueva and Chen, 2019). Analysis scripts are provided in Supplemental File 1 and data are provided in Supplemental File 2. The observed annual precipitation, averaged across locations, was used to classify years into three “precipitation level” quantiles: “dry” years (161–428 mm), “moderate” years (428–580 mm), and “wet” years (580–1380 mm). Note, this classification is not intended as a formal envirotyping of water deficit scenarios (Chenu et al., 2013) but to reflect the type of informal classification used by growers and agronomic practitioners in the region (Baeumler and Gupta, 2020). The effects of planting date and the interaction of planting date and precipitation levels on all variables were tested using analysis of variance (ANOVA), considering planting date, precipitation level, and hybrid type as fixed factors, and location and year as random factors. Tukey test for the interaction of planting date and precipitation levels was performed when the *F* value was below an α < 0.05 significance threshold. Daily transpiration and water stress events were assigned to vegetative and reproductive stages. Daily soil evaporation was aggregated into fallow and growing periods. Soil evaporation was identical from January 1 to April 15 across all three planting dates, and not modeled from the date of the latest harvest (typically October) to December 31, so these periods were ignored.

## RESULTS

### Model performance for days to emergence

The CERES-Sorghum model performance for days to emergence using a Tbase of 6 °C, 8 °C, and 10 °C was evaluated by comparing observed and simulated days to emergence for experiments conducted on different planting dates in Kansas. Results indicated a RMSE of 2.2, 1.7, and 2.0 days for a Tbase of 6 °C, 8 °C and 10 °C, respectively (Figure 2a). For six experiments planted in April, observed days to seedling emergence ranged from 13 to 19 days, while simulated days to emergence, with a Tbase of 8 °C, ranged from 11 to 18 days (Figure 2a). For two experiments planted in May, observed days to emergence were 5 and 7 and the corresponding simulated values, with a Tbase of 8 °C, were 6 and 8 days, respectively (Figure 2a).

Seasonal simulations for days to emergence for a Tbase ranging from 6 °C to 10 °C in the study site (Figure 2b) indicated that, for instance, in northwest location (Colby) an increase in 1 °C for Tbase delays the crop emergence around 1.5, 0.6, and 0.2 days for planting dates in mid-April, mid-May and mid-June, respectively. Across locations, an increase in 1 °C delays the crop emergence in around 1.3, 0.5, and 0.2 for planting dates in very early, early, and normal planting dates, respectively. Overall, these results suggest that a lower Tbase reduces the time required for emergence, particularly when the crop is simulated to be planted in early spring (April 15).

Therefore, as outlined above, simulations in early spring using a Tbase of 8 °C do represent days to emergence for kaoliang, a chilling tolerant genotype, when planted in early spring.

### Soil moisture at planting

At each location and year, simulations started on January 1 to accrue soil moisture in the soil profile as a function of daily precipitation (Figure 3). In this growing region, high variability of monthly precipitation is observed from May to August (Figure 3a). The direct impact of precipitation in soil moisture is illustrated in Figure 3b and 3c, where the soil moisture or extractable soil water on each planting date varied across locations, with less soil moisture in western locations than in eastern locations and less soil moisture in very early and early planting (April 15, May 15) dates than in normal planting dates (June 15). In general, less moisture accrued prior to planting for very early planting dates (158 ∓35 mm) compared to early (168 ∓40 mm) and normal planting dates (185 ∓46 mm).

**Figure 3.**
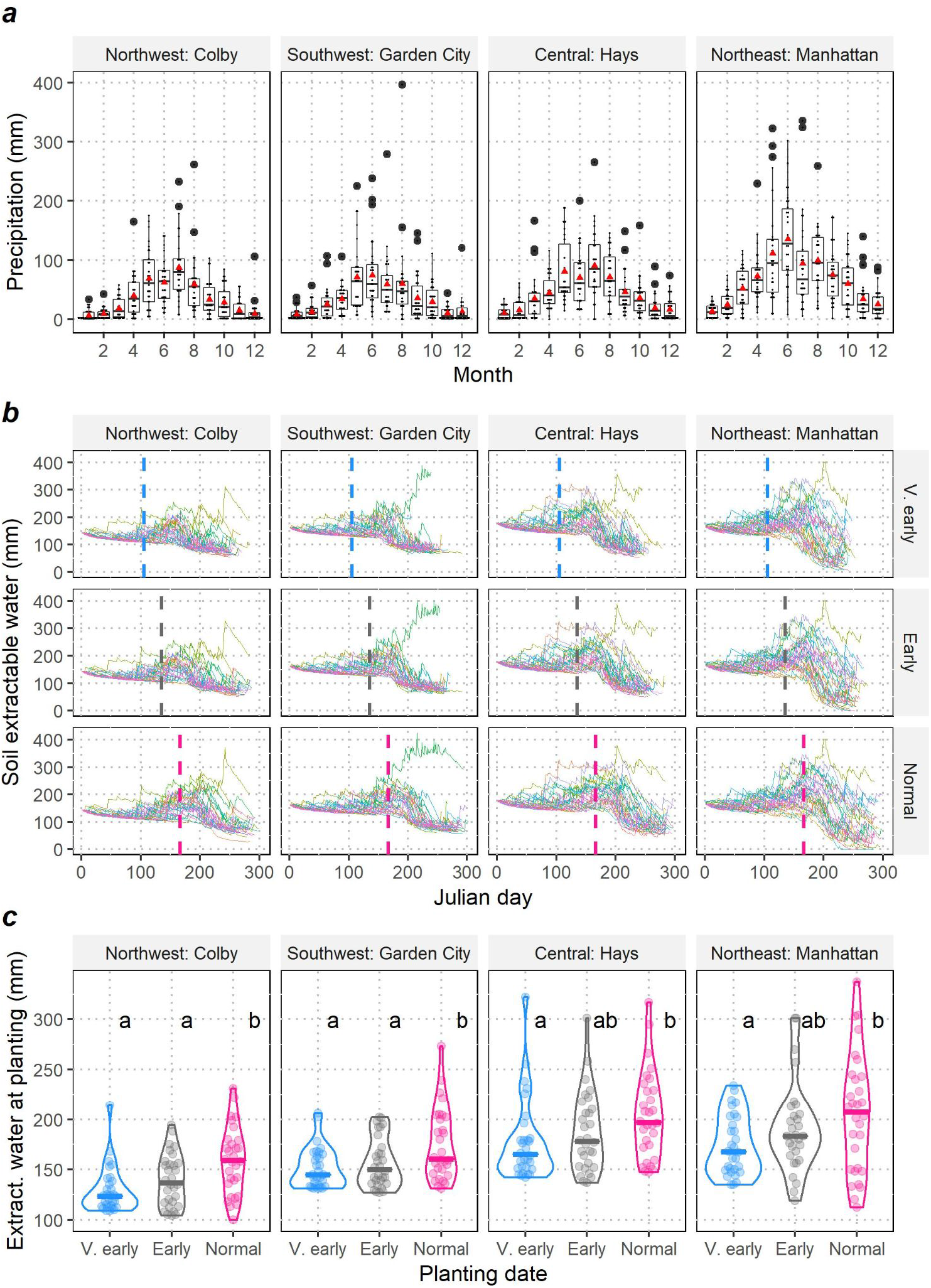
Effect of precipitation on soil moisture at planting for a simulated full-season hybrid. (a) Variability of monthly precipitation in four representative locations in Kansas, red triangles represent the monthly mean over a 30 years period. (b) Daily soil extractable water (soil moisture) in different planting dates, vertical dashed lines indicate the planting date at each location. (c) Extractable water for very early (mid-April), early (mid-May), and normal (mid-June) dates planting. Each boxplot and violin plot represent the annual variability over 30 years (1986-2015) in Kansas. Letters represent significant differences (α < 0.05) using the Tukey HSD test.

### Growing season length

In each year, the phenology development and corresponding environmental conditions were classified into days from planting to seedling emergence, vegetative growth (days from seedling emergence to anthesis), and reproductive growth (days from anthesis to physiological maturity) (Figure 4, Table S2). Simulations indicated that shifting planting dates from early summer to early spring slowed down the rate of seedling emergence. For instance, in very early, early, and normal planting dates seedlings are expected to emerge 16, 10, and 6 days after planting, respectively. The number of days to seedling emergence and vegetative growth were greater in western regions than in eastern regions (Figure 4a and 4b). For both full- and short-season hybrids, very early planting dates extended the vegetative stage and shortened the reproductive stage. By contrast, normal planting dates shortened the vegetative stage and extended the reproductive stage (Figure 4, Table S2). For instance, a full-season hybrid that was simulated to be planted in early spring (April 15) and early summer (June 15) completed the vegetative growth in 79 and 63 days and reproductive growth in 44 and 48 days, respectively (Table S2).

**Figure 4.**
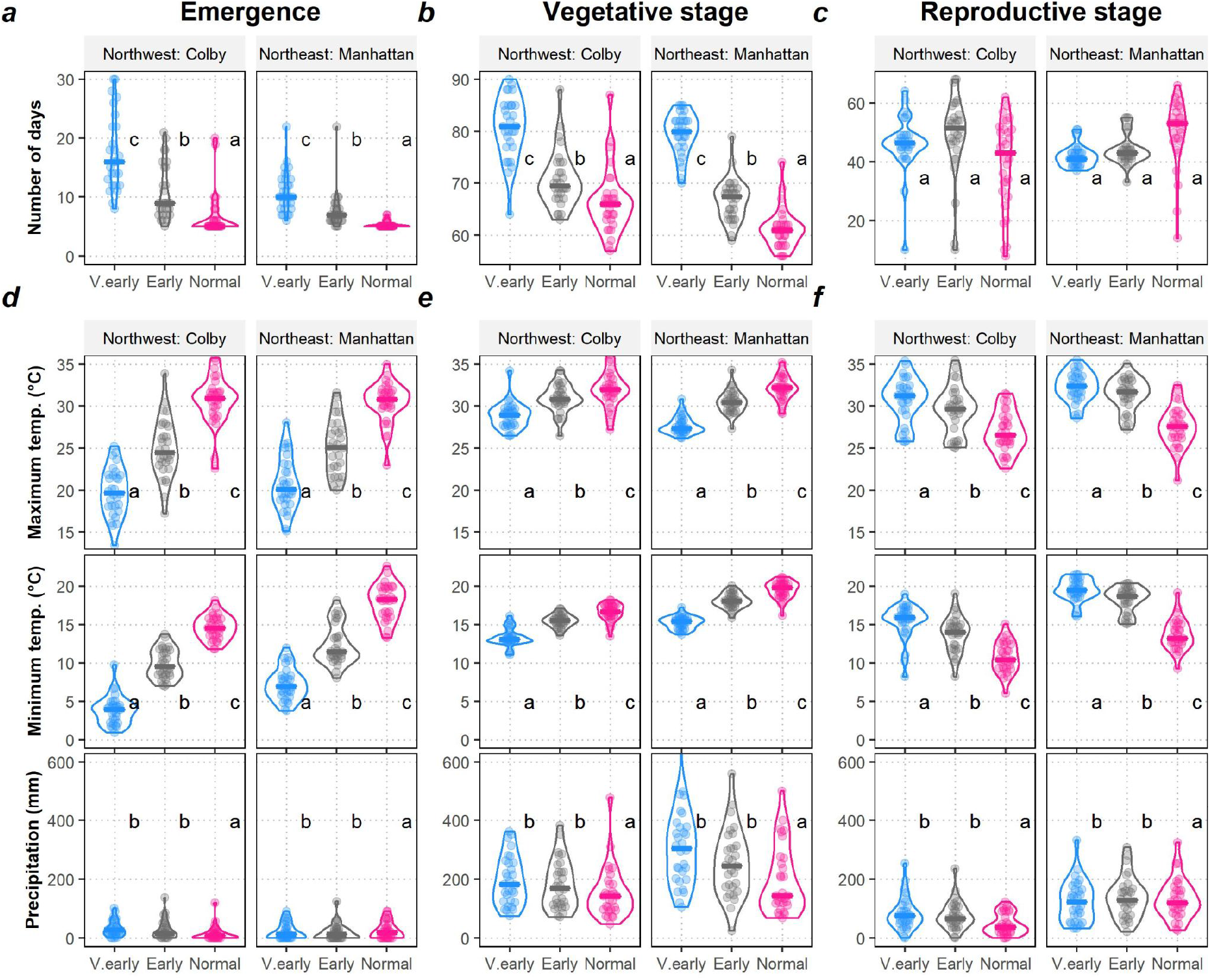
Crop phenology for a simulated full-season sorghum hybrid in two contrasting locations. (a) Days for emergence, (b) days for vegetative development and (c) days for reproductive development for very early (mid-April), early (mid-May), and normal (mid-June) planting dates. Environmental conditions during (d) days to emergence, (e) days for vegetative, and (f) reproductive stage in different planting dates. Each violin plot represents the annual variability of 30 years (1986-2015) at each location in Kansas. Letters indicate significant differences (α < 0.05) using the Tukey HSD test.

### Temperatures experienced by growth stage

Very early, early and normal planting scenarios led to contrasting temperature profiles during the growing period (Figure 4d-f), as summarized by the maximum (Tmax) and minimum (Tmin) temperatures experienced by the crop at the given stage (Table S2). When a full-season hybrid was simulated to be planted very early (mid-April), it experienced a temperature trend from low (vegetative stage) to high (reproductive stage). By contrast, when planted in mid-June, it experienced a temperature trend from high (vegetative stage) to low (reproductive stage).

In general, across all three planting dates most of the precipitation (from 63–65%) occurred during the vegetative growth (Table S2). On average, in very early planting dates the crop experienced slightly higher seasonal precipitation compared to normal planting (347 mm vs. 263 mm for full-season hybrids; 310 mm vs. 240 mm for short-season hybrids). In western sites, temperatures were lower than in the eastern sites. For instance, given normal planting, during reproductive stages the minimum temperature in the northwestern site (Colby) was 11 °C and in the eastern site (Manhattan) was 14 °C.

### Soil evaporation in fallow and growing periods

Daily soil evaporation was aggregated into fallow and growing periods and compared across planting date and hybrid type. Simulations of a full-season hybrid indicate that on average across all planting dates, 30% of soil evaporation occurred in fallow periods and 70% during growing periods. Most of the soil evaporation during the growing period occurred before anthesis (53% of total soil evaporation). For all precipitation levels (dry, moderate, and wet years), very early planting dates significantly reduced the total soil evaporation compared with normal planting dates (Figure 5a, upper panel). Likewise, for all precipitation levels, very early planting dates reduced the soil evaporation during fallow periods (Figure 5a, lower panel, upper bars light colors) but increased the soil evaporation during the growing periods (Figure 5a, lower panel, lower bars and dark colors).

**Figure 5.**
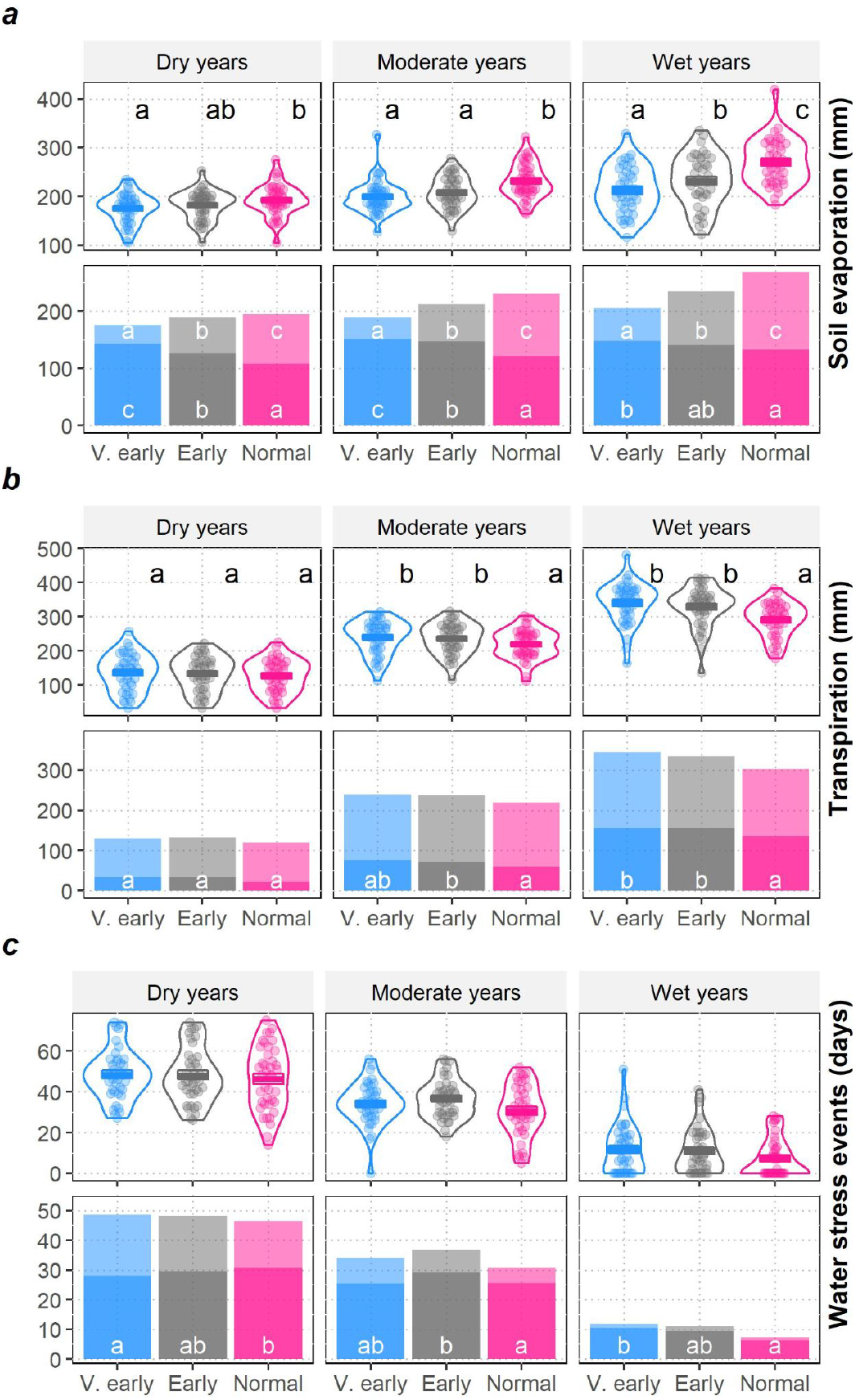
Water budgets under different levels of precipitation for a simulated full-season sorghum hybrid. (a) Total soil evaporation across planting dates (upper panel), soil evaporation during fallow periods (upper bars, light colors) and growing periods (lower bars, dark colors) under different planting dates. (b) Plant transpiration or water capture during the growing period (upper panel), and during vegetative (upper bars, light colors) and reproductive stages (lower bars, dark colors) under different planting dates. (c) Water stress events during the growing period (upper panel), and during vegetative (upper bars, light colors) and reproductive stages (lower bars, dark colors). Each violin plot represents the annual variability of 30 years (1986-2015) in Kansas. Each bar plot represents the median of each variable on each phenological stage. Letters represent significant differences (α < 0.05) using the Tukey HSD test. Annual precipitation for dry, moderate, and wet years was 161-428 mm, 428-580 mm and 580-1380 mm, respectively.

### Plant transpiration

Daily transpiration for simulated full- and short-season hybrids was aggregated into vegetative and reproductive growth for each planting date in each year (Table S3). As would be expected, transpiration was substantially higher in the wet years (289 mm) compared to moderate (212 mm) or dry years (128 mm) across all planting dates. Simulations for a full-season hybrid indicated higher average total transpiration or water capture for very early planting dates (236 mm) than early (232 mm) or normal planting dates (211 mm). In dry years, total transpiration was similar in all planting dates (Figure 5b, upper panel). In both, moderate and wet years, very early and early planting dates significantly increased plant transpiration compared with late planting dates (Figure 5b, upper panel).

During vegetative growth, no significant differences in transpiration were found among planting dates in all precipitation levels (Figure 5b, lower panel, upper bars and light color). During reproductive growth, discrepancies in transpiration among planting dates were negligible only in dry years. However, these discrepancies in transpiration among planting dates were consistent, in both, moderate and wet years (Figure 5b, lower panel, lower bars and dark colors). Overall for a full-season hybrid, in dry years, transpiration during reproductive growth in very early and normal planting dates averaged 27% and 21%, respectively. By contrast, in wet years, transpiration during reproductive growth was greater, averaging 44% and 42% of transpiration for very early and normal planting, respectively. This suggests that very early and early planting can lead to drought escape under favorable scenarios.

### Water stress events

CERES-Sorghum has two water stress factors whose values range from 0 (no stress) to 1 (stress). In this study a water stress event is defined when the minimum value between water deficit factor for photosynthesis (SWFAC) or water deficit factor for development (TURFAC) was higher than 0.6. Next, the number of water stress events were aggregated into vegetative and reproductive growth and are presented in Table S4. Simulations for a full-season hybrid indicate that on average the number of days under water stress during the growing period in very early, early, and normal planting dates was 31, 32 and 28, respectively. Similarly, on average across all planting dates the number of water stress events during the growing period was higher in dry years (39), followed by moderate (30) and wet years (12).

The number of days under water stress in different planting dates and precipitation levels during the growing period for a full-season hybrid is provided in Figure 5c. During the growing period (Figure 5c, upper panel) and during vegetative stage (Figure 5c, upper bars and light color) the frequency of water stress events among planting dates was negligible in all precipitation levels. During the reproductive stage the frequency of water stress events was significantly higher in very early and early planting dates in all precipitation levels (Figure 5c, lower bars and dark color).

### Grain yield

CERES-Sorghum simulates the plant-soil-water dynamics on a daily time step that finally is translated into grain yield. To evaluate the effect of planting date on grain yield, we have conducted simulations for full- and short-season hybrids (Figure 6 and Table S5). The simulated grain yield over 1986–2015 averaged 4 Mg ha^−1^ across locations, planting dates, and hybrids, and ranged from 0 to 12 Mg ha^−1^. Zero yield was obtained in 1989 in the northwest site (Colby) for all planting dates due to exceptionally low annual precipitation (161 mm) and in 2004 in the southwest site (Garden City) due to low precipitation in very early spring. In general, grain yield varied significantly among planting dates (*p* < 0.001), precipitation levels (*p* < 0.001) and hybrids (*p* < 0.001).

**Figure 6.**
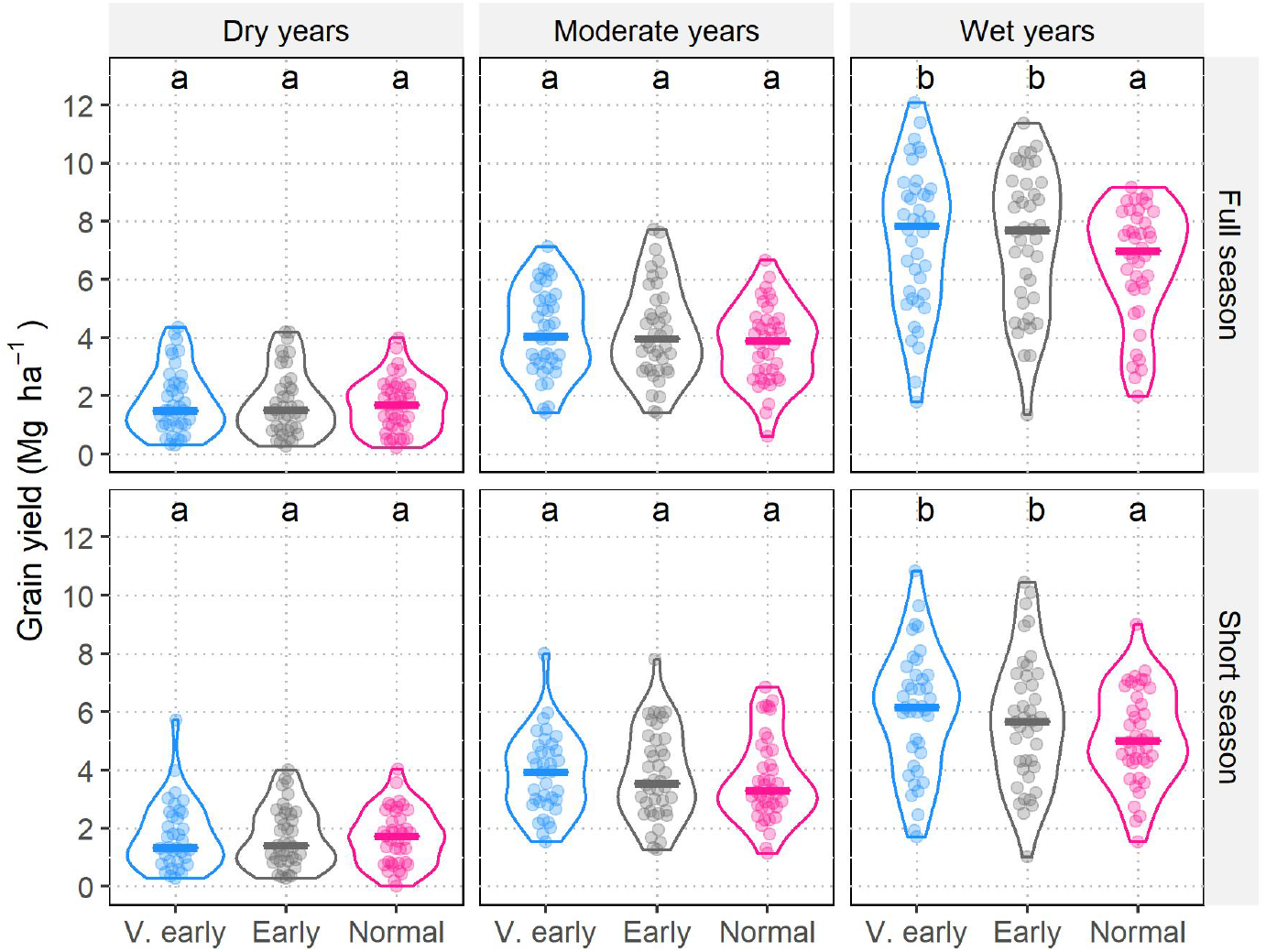
Grain yield under different levels of precipitation and planting dates for simulated full- and short-season hybrids. Each violin plot represents the annual variability of 30 years (1986-2015) in Kansas. Letters indicate significant differences (α < 0.05) of all pairwise comparisons using the Tukey HSD test. Annual precipitation for dry, moderate, and wet years was 161-428 mm, 428-580 mm, and 580-1380 mm, respectively.

Low grain yield was obtained in dry years, as expected, and differences in grain yield among planting dates were negligible. For instance, in dry years, the simulated yield for full-season hybrid was 1.6 Mg ha^−1^, not significantly different (*p* > 0.05) from yields of either very early, early or normal planting dates. In wet years, by contrast, yields were higher for very early and early planting versus normal planting. For instance, in wet years very early planting dates averaged 7.4 Mg ha^−1^ for a full-season hybrid, 13% greater than the mean for normal planting dates (6.5 Mg ha^−1^).

Overall, interactions between planting dates and hybrids were not significant (Table S5), indicating similar trends for both maturing genotypes with very early and early planting dates with yield increase in environments from moderate to high precipitation. Nevertheless, grain yield across locations for full- and short-season hybrids in contrasting sites (Table S6) indicated that short-season hybrids can contribute a slight yield increase for normal planting dates. For instance, in the northwestern site (Colby) under normal planting, the grain yield of a short-season hybrid was 2.9 Mg ha^−1^ and for a full-season hybrid was 2.6 Mg ha^−1^. Overall, results indicate that very early and early planting of chilling-tolerant sorghum hybrids can potentially increase grain yield in 66% of the years.

## DISCUSSION

### Validity of the modeling approach for the target cropping system

After more than 50 years of breeding for the early-planted chilling tolerance trait in grain sorghum, this study provides insights on the impact of this G × E × M intervention at crop scale, narrowing the bridge between genotype to phenotype. To this end, this study used an agricultural systems tool, the CERES-Sorghum model, because it is one of the most widely used sorghum models (Adam et al., 2018; Amouzou et al., 2019; Folliard et al., 2004; Fu et al., 2016; Kothari et al., 2020; Pachta, 2007; Singh et al., 2014), and its accuracy has been established for simulations of grain yield in Kansas for normal planting dates (Araya et al., 2017; Staggenborg and Vanderlip, 2005; White et al., 2015). Further, the model accuracy in predicting days of emergence for early and normal planting dates (Figure 2a) and the sensitivity analysis for Tbase (Figure 2b) suggests that the CERES-Sorghum model, which uses a Tbase of 8 °C, is likely appropriate to forecast the effects of variables analyzed in this study.

Still, the congruence between observed and simulated values for soil moisture, evapotranspiration, soil evaporation, and transpiration for the CERES-Sorghum model has not yet been demonstrated. Crop model applications rely on rigorous model testing or model comparison between observed versus simulated variables (Wallach et al., 2014). Other models such as APSIM-Maize and APSIM-Soybean simulated field observations of soil moisture for maize and soybean (Ebrahimi-Mollabashi et al., 2019; Yang et al., 2018). Likewise, observed seasonal evapotranspiration for sweet sorghum and maize matched those values simulated with crop models CERES-Sorghum and CERES-Maize (Araya et al., 2017; DeJonge et al., 2012; Lopez et al., 2017). Commonly, in sorghum and other cereal crops, crop model evaluation is conducted for grain yield and biomass components (Kassie et al., 2016; Yakoub et al., 2017). Consequently, field experimentation to evaluate water budgets are needed for grain sorghum.

Some aspects of farm conditions not considered by the model may affect the interpretation of our findings. For instance, simulated fallow periods before planting disregard weed growth pressure that can deplete soil moisture and nutrients. Studies in Kansas indicate that in dry years, green fallow or a crop can deplete soil moisture decreasing the productivity of the subsequent winter wheat season (Holman et al., 2016; Schlegel et al., 2017; Schlegel and Havlin, 1997). Note, the fallow period before planting, which allows weed growth, was longer for normal planting dates versus the earlier planting dates. For this reason, it is possible that the model overestimates expected soil moisture available at this planting date.

### Potential benefits of early-planted chilling tolerant sorghum

In the US maize and sorghum belts, optimum planting windows for maize and soybean vary between mid-April to late-May (Baum et al., 2019; Long et al., 2017; Mourtzinis et al., 2017; Staggenborg et al., 1999a); therefore, these optimum planting windows would be suitable for sorghum. Indeed, simulations show that bringing forward the sorghum planting date in mid-April (very early) and mid-May (early) in years with moderate to high precipitation (top two-thirds of years based on annual precipitation) can lead to a yield increase on average by 12% for a full-season hybrid and and 7% for a short-season hybrid compared to normal planting dates (Table S5). Higher sorghum yields were reported in Kansas in planting dates in early June (Ciampitti et al., 2019). By contrast, when shifting planting dates from June to May, a yield increase (4–35%) was reported in four experiments while a yield decline (−69%) was reported in one experiment (Maiga, 2012). Those experimental findings are in line with our simulations and indicate that early chilling tolerance can increase yields under favorable scenarios (Figure 6). Given that yields in very early and early planting dates were not substantially different (Figure 6, Table S5), it is likely that the optimum planting window for chilling tolerant sorghum assuming an adequate early stand establishment ranges from mid-April to mid-May, similar to those of maize and soybean.

In cereals, drought escape can be achieved through short-season varieties that flower earlier (Blum, 2011; Shavrukov et al., 2017), and in sorghum drought escape is a proposed benefit of early planting and chilling tolerance (Burow et al., 2011; Franks et al., 2006). However, our simulations suggest that the probability of water stress during grain filling would actually be expected to increase slightly for planting dates in early spring (Figure 5c). The results to some extent support that it is possible to avoid the terminal drought by increasing plant transpiration or water capture in years with moderate to high precipitation (Figure 5b). In our simulations, the reduction of soil evaporation during the fallow portion of the season can not directly be capitalized into grain yield, but this water budget can hypothetically benefit a double crop rotation (Burow et al., 2011) or be available in the soil profile for the subsequent cropping season (Figure S2). In dryland systems of western Kansas, wheat-sorghum is a common crop rotation due to benefits in grain yield and soil water moisture (Schlegel and Havlin, 1997). Using an early-planted chilling-tolerant grain sorghum in this rotation can potentially further improve grain yield and water productivity for both crops. Additional simulations of crop rotations would be needed to assess these benefits.

### Expected tradeoffs for early and normal planting dates

Environmental conditions dictate the rate of phenology development. Planting dates in early spring extended the duration of the growing season (Figure 4, Table S2), especially days for seedling emergence and days to anthesis, as previously predicted (Burow et al., 2011) and observed (Kapanigowda et al., 2013). The delay in seedling emergence in very early spring (Figure 4) was additionally associated with low levels of soil moisture (Figure 3). Although, studies on chilling tolerance in field conditions (Maulana and Tesso, 2013; Ostmeyer et al., 2020) do not report the effect of soil moisture on germination and seedling emergence. Soil moisture deficit and low temperature both impair germination and delay emergence in sorghum (Evans et al., 1961), but their interactions are not well understood. Consequently, our study suggests that hybrids with chilling tolerance should be planted under optimum levels of initial soil moisture to ensure early crop establishment, as recommended for other crops such as maize (Lu et al., 2017).

Sorghum grain yield is affected when the crop experiences extreme temperatures, above 38 °C and below 11 °C, during grain filling (Singh et al., 2015; Staggenborg et al., 1999b). In our simulations, very early planting exposed the crop to moderately high temperatures (<38 °C) during grain filling. By contrast, normal planting exposed the crop to temperatures below 10 °C during grain filling in 50% of the years in the northwest site (Colby) (Figure 4f and Table S6). Risk of freezing temperatures for normal planting dates were reported in studies conducted in Kansas, Colorado and Texas (Baumhardt and Howell, 2006; McMaster et al., 2016; Staggenborg et al., 1999b; Staggenborg and Vanderlip, 1996). Furthermore, in Colorado short-season hybrids can avoid freezing temperatures during grain filling for planting dates in late spring (McMaster et al., 2016).

Climate change, which was not accounted for in our simulations, may affect the potential benefits and tradeoffs for very early and early planting of chilling-tolerant sorghum. For instance, climate time series analysis in Kansas indicates that the frost risk events have become less frequent in the last decade (Lin et al., 2017) and global temperatures are expected to rise another 0.5–2.5 °C by mid-century (IPCC, 2013). In future scenarios it is possible that the risk of heat stress during flowering can increase for very early planting, while the frequency of frost damage may not be important for planting dates in early summer. Alternatively, very early planting of chilling-tolerant hybrids could provide heat stress escape. Simulations under future climate scenarios would be needed to resolve these potential tradeoffs.

A potential risk to very early, early and normal plantings was the low grain yield in years with low precipitation (Figure 6), due to the low soil moisture (Figure 3) and high frequency of daily droughts during pre-flowering and post-flowering stages (Figure 5c). This effect was reported in other crops. For instance, in Iowa, locations exposed to earlier water stress exhibited lower soybean yield (Irmak et al., 2002), while across the US midwest, locations with low soil moisture imposed a greater penalty on maize yield (Seifert et al., 2017). In Texas, a simulation study indicated that a short-season hybrid is better adapted to dryland systems when the crop is planted in June (Baumhardt and Howell, 2006). Similar results were obtained for the northwestern site (Colby), where a short-season hybrid yielded higher than a full-season hybrid in planting dates in late spring (Table S6). Overall, low early-season water budgets as a result of low precipitation can override the value of very early planting.

### Next steps for models of chilling tolerance to guide crop improvement

In breeding and genetics studies, chilling tolerance is most often characterized through seedling emergence rate and seedling vigor scores, usually based on visual evaluation (Burow et al., 2011; Franks et al., 2006; Marla et al., 2019; Parra-Londono et al., 2018). By contrast, crop models simulate days to emergence, seedling biomass weight, and leaf area. CERES-Sorghum, developed for normal conditions, does not model any detrimental effect of chilling temperatures with respect to seedling emergence or leaf damage. Consequently, the model can not simulate the grain yield for a genotype which was damaged by chilling temperatures.

Future work to extend a chilling sensitive routine could entail penalizing the development rate, growth rate, or leaf area. The development rate can be reduced by shifting the base temperature (e.g. from 8 °C to 10 °C) and varying the duration from planting to emergence. For instance, currently thermal time from planting to germination (50 degree days) and the coleoptile extension rate (0.1 °C day cm^−1^) are fixed parameters. Chilling tolerant genotypes can attain 50% seedling emergence two days earlier than their susceptible counterparts (Stickler et al., 1962), suggesting that there is variability for base temperature, thermal time from planting to germination, and/or coleoptile extension rate. Otherwise, differences for base temperature (0–9.8 °C) were recently reported for Ethiopian sorghum germplasm, which is believed to harbor adaptations to chilling-prone high-altitude regions (Tirfessa et al., 2020).

In this study, simulations were aggregated into three precipitation levels using a simple quantile classification. Although the frequency of water stress corresponded well to each precipitation level (Figure 5c) the daily water stress can be used to characterize drought patterns more precisely. For instance, the APSIM-Wheat and the SSM model were used to classify and quantify drought patterns across the Australian wheat belt (Chenu et al., 2013) and the US maize belt (Messina et al., 2015), respectively. Thus, the characterization of drought patterns (i.e. envirotyping) with CERES-Sorghum, in these locations and across the US sorghum belt would guide crop improvement programs to focus on traits better-suited for target environments.

Overall, through years of field experimentation, research on early season chilling tolerance pointed out the potential outcomes of a sorghum hybrid with this trait. This study has shown that a crop simulation model can translate the genetic value of this breeding trait in terms of grain yield by integrating environmental conditions (i.e. the impact of annual variability) and realistic agronomic management. Furthermore, this study tested hypotheses regarding benefits of early chilling tolerance proving that it is difficult to informally predict the state of any variable (i.e. evapotranspiration) at crop system scale, without formal modeling. Thus, our study underscores the value of crop modeling to complement and guide plant breeding and genetics.

## ACKNOWLEDGEMENTS

This study was supported by funding from the Foundation for Food and Agriculture Research - Seeding Solution “CA18-SS-0000000094 – Bridging the Genome-to-Phenome Breeding Gap for Water-Efficient Crop Yields (G2P Bridge)” to G.P.M. and I.C; the Kansas Department of Agriculture “Collaborative Sorghum Investment Program Water Optimized Sorghum for Kansas” to G.P.M; and the Kansas Grain Sorghum Commission.

## AUTHOR CONTRIBUTIONS

Designed the research: GPM, RR, IC. Performed research: RR. Analyzed data: RR, SSB. Wrote the paper: RR, GPM.

## SUPPLEMENTARY DATA

### Included in the review PDF file

Supplementary Tables 1–6

Supplementary Figures 1–2

### Uploaded separately

Supplementary File 1 - Code

Supplementary File 2 - Data

---

**Table S1.** Soil physical characteristics of study sites in Kansas.

| Location | Depth (cm) | LL (cm cm<sup>-1</sup>) | DUL (cm cm<sup>-1</sup>) | SAT (cm cm<sup>-1</sup>) | RGF (-) | BD (g cm<sup>-3</sup>) | OC (%) | Clay (%) | Silt (%) | pH |
| --- | --- | --- | --- | --- | --- | --- | --- | --- | --- | --- |
| Colby | 0-10 | 0.21 | 0.39 | 0.43 | 1.00 | 1.43 | 1.16 | 30 | 52 | 7.2 |
|  | 10-27 | 0.25 | 0.43 | 0.43 | 1.00 | 1.43 | 0.87 | 39 | 54 | 7.5 |
|  | 10-56 | 0.16 | 0.34 | 0.44 | 0.44 | 1.43 | 0.29 | 25 | 63 | 7.5 |
|  | 56-194 | 0.16 | 0.32 | 0.49 | 0.08 | 1.28 | 0.29 | 25 | 52 | 8.5 |
| Garden City | 0-15 | 0.18 | 0.37 | 0.49 | 1.00 | 1.3 | 1.2 | 24 | 60 | 7.9 |
|  | 10-32 | 0.17 | 0.35 | 0.48 | 0.63 | 1.3 | 0.6 | 26 | 59 | 7.9 |
|  | 32-74 | 0.15 | 0.33 | 0.46 | 0.35 | 1.4 | 0.3 | 23 | 64 | 8.5 |
|  | 74-194 | 0.15 | 0.33 | 0.46 | 0.07 | 1.4 | 0.3 | 23 | 64 | 8.5 |
| Hays | 0-15 | 0.18 | 0.39 | 0.43 | 1.00 | 1.44 | 1.16 | 24 | 67 | 6.7 |
|  | 15-22 | 0.19 | 0.40 | 0.47 | 1.00 | 1.31 | 1.16 | 26 | 67 | 6.7 |
|  | 22-32 | 0.22 | 0.40 | 0.45 | 0.58 | 1.38 | 0.58 | 35 | 60 | 7.3 |
|  | 32-64 | 0.23 | 0.41 | 0.45 | 0.38 | 1.38 | 0.46 | 39 | 56 | 7.3 |
|  | 64-88 | 0.18 | 0.37 | 0.44 | 0.22 | 1.42 | 0.29 | 30 | 64 | 7.9 |
|  | 88-194 | 0.16 | 0.35 | 0.44 | 0.06 | 1.42 | 0.29 | 25 | 69 | 7.9 |
| Manhattan | 0-17 | 0.19 | 0.41 | 0.44 | 1.00 | 1.4 | 1.7 | 24 | 64 | 6.5 |
|  | 17-29 | 0.21 | 0.40 | 0.45 | 0.63 | 1.4 | 1.2 | 30 | 58 | 6.5 |
|  | 29-39 | 0.22 | 0.40 | 0.44 | 0.51 | 1.4 | 0.9 | 33 | 58 | 6.5 |
|  | 39-69 | 0.25 | 0.43 | 0.44 | 0.34 | 1.4 | 0.6 | 42 | 51 | 6.7 |
|  | 69-91 | 0.23 | 0.40 | 0.45 | 0.20 | 1.4 | 0.4 | 39 | 54 | 6.7 |
|  | 91-130 | 0.21 | 0.38 | 0.44 | 0.11 | 1.4 | 0.2 | 36 | 54 | 7.2 |
|  | 130-194 | 0.19 | 0.36 | 0.44 | 0.04 | 1.4 | 0.1 | 32 | 56 | 7.2 |

LL, DUL, SAT and RGF were estimated using a pedotransfer function embedded in the Sbuild tool of DSSAT suit.
LL: lower limit
DUL: drainage upper limit
SAT: saturation
RGF: root growth factor
BD: bulk density
OC: organic carbon

**Table S2.**
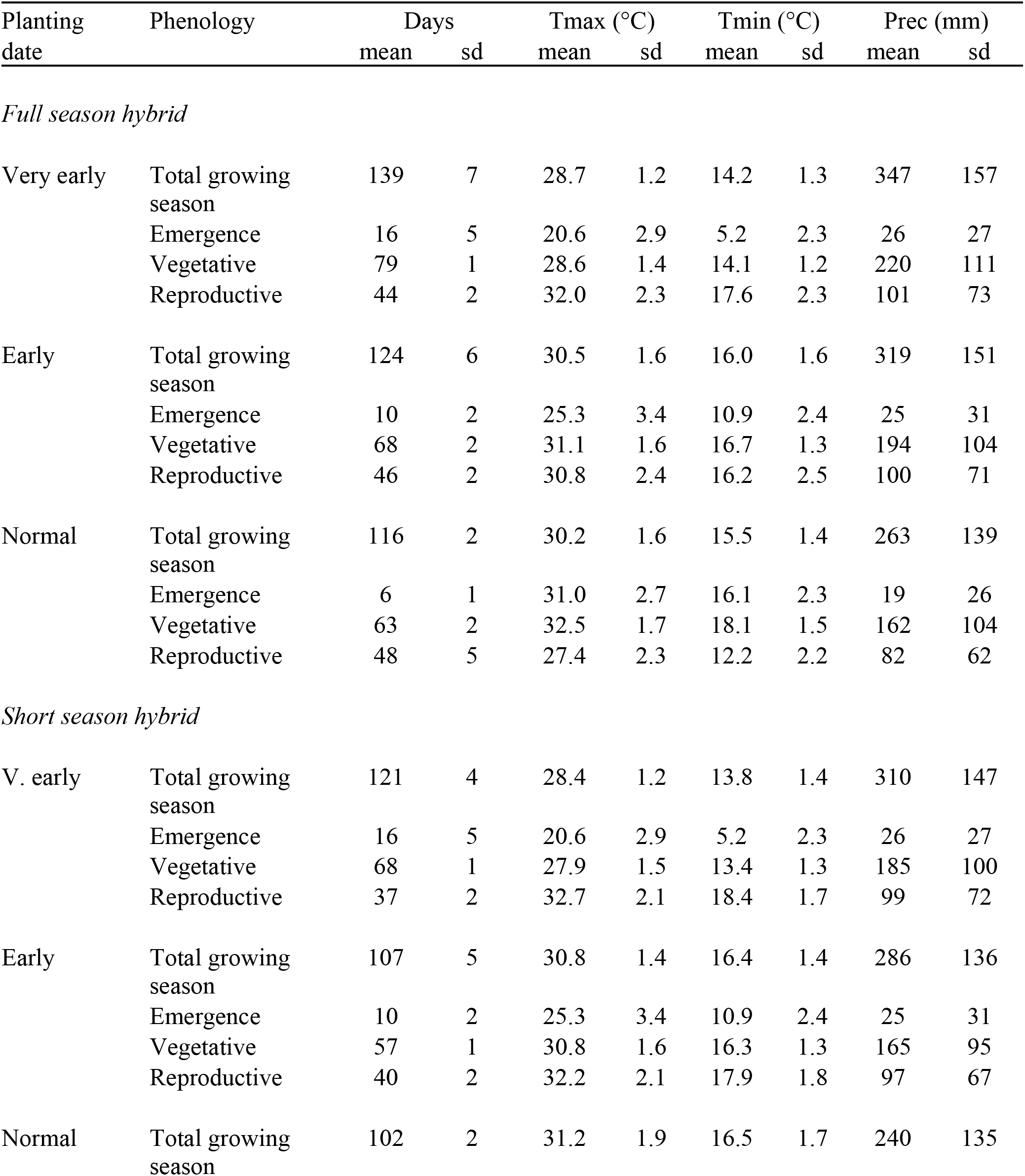
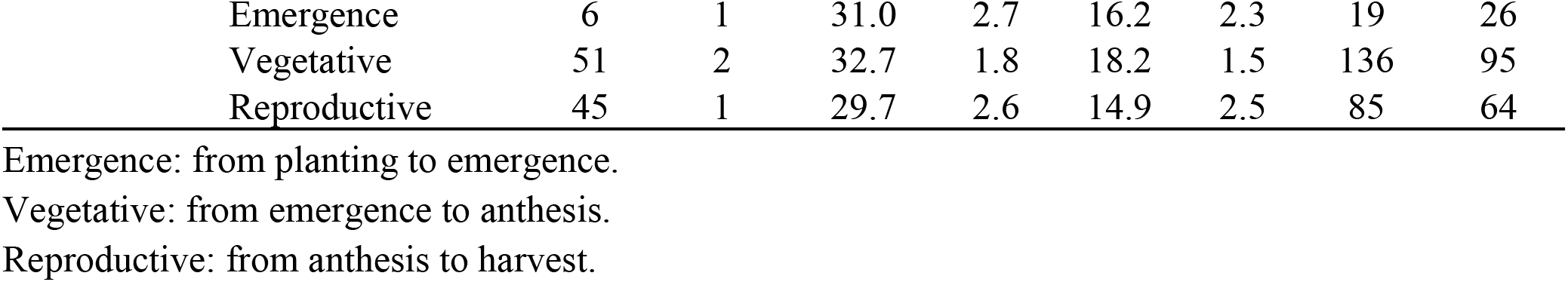
Phenology development and environmental conditions during the growing period and during vegetative and reproductive stage for a simulated full- and short-season hybrids in three planting dates across representative sites in Kansas. Each value represents the overall mean over four locations and 30 years (1986-2015).

**Table S3.** Transpiration (mm) during the growing period and during vegetative and reproductive stage for a simulated full- and short-season hybrids in three planting dates across representative sites in Kansas. Each value represents the great mean over four locations and 30 years (1986-2015).

| Hybrid | Planting date | Total<br>(mm) |  | Vegetative<br>(mm) |  | Reproductive<br>(mm) |  |
| --- | --- | --- | --- | --- | --- | --- | --- |
|  |  | mean | sd | mean | sd | mean | sd |
| Full season |  |  |  |  |  |  |  |
|  | V. early | 236 | 104 | 149 | 58 | 87 | 61 |
|  | Early | 232 | 97 | 147 | 53 | 86 | 58 |
|  | Normal | 211 | 84 | 141 | 44 | 70 | 53 |
| Short season |  |  |  |  |  |  |  |
|  | V. early | 200 | 99 | 119 | 56 | 80 | 56 |
|  | Early | 202 | 81 | 120 | 42 | 81 | 52 |
|  | Normal | 178 | 67 | 107 | 36 | 72 | 44 |

| Source of variation | df | Total | Vegetative | Reproductive |
| --- | --- | --- | --- | --- |
| Planting date (PD) | 2 | <0.001 | <0.001 | <0.001 |
| Hybrid (V) | 1 | <0.001 | <0.001 | 0.1 |
| Precipitation levels (dnw) | 2 | <0.001 | <0.001 | <0.001 |
| PD x dnw | 4 | <0.001 | 0.3 | <0.001 |

**Table S4.**
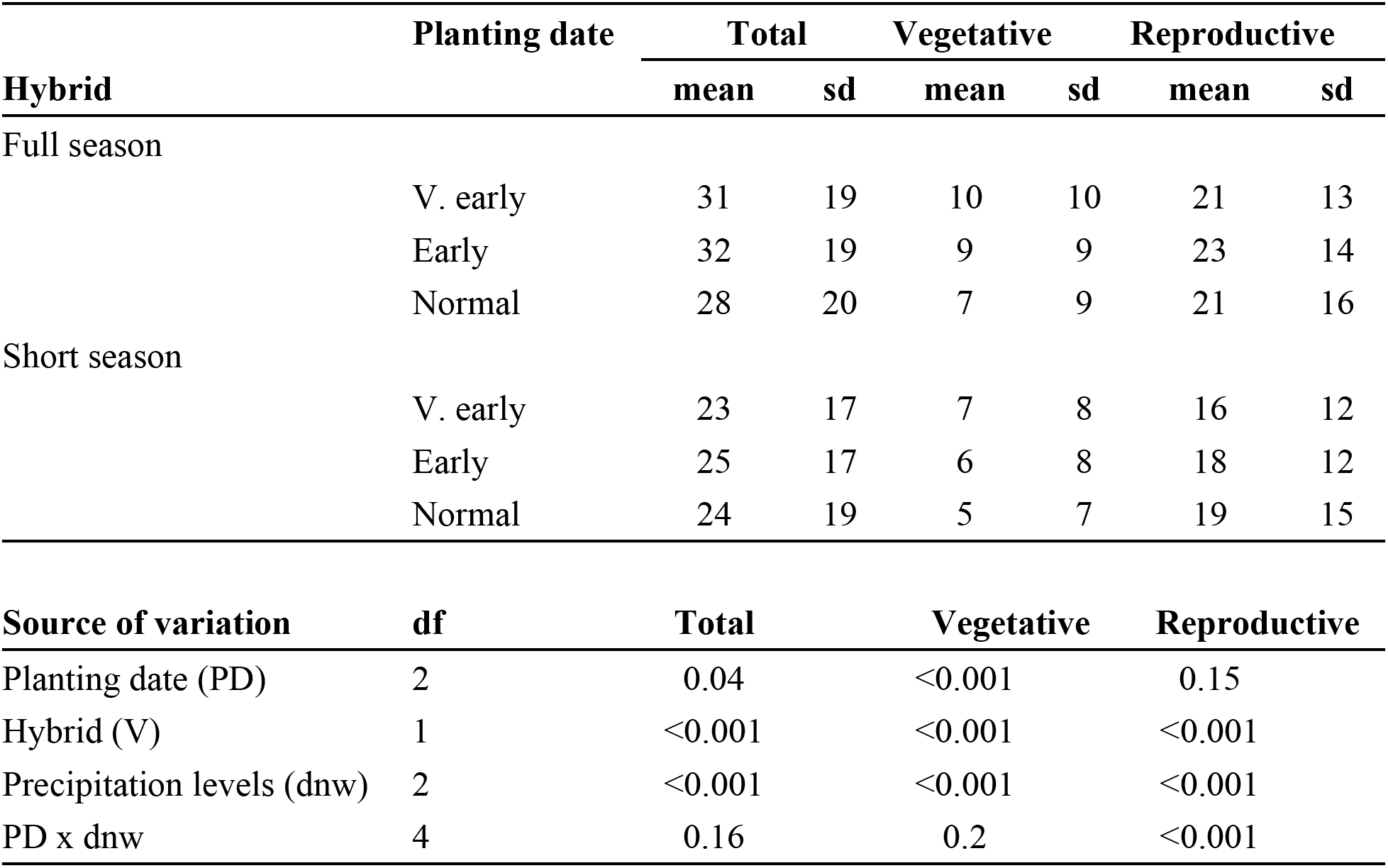
Water stress events (days) during the growing period and during vegetative and reproductive stage for a simulated full- and short-season hybrids in three planting dates across representative sites in Kansas. Each value represents the great mean over four locations and 30 years (1986-2015).

**Table S5.** Grain yield (Mg ha^−1^) for a simulated full- and short-season hybrids in three planting dates across representative sites in Kansas. Each value represents the great mean over four locations and 30 years (1986-2015).

| Hybrid | Planting date | Grain (Mg ha<sup>-1</sup>) |  | group (PDxV) |
| --- | --- | --- | --- | --- |
|  |  | mean | sd |  |
| Full season | V. early | 4.4 | 2.9 | b |
|  | Early | 4.4 | 2.9 | b |
|  | Normal | 4.0 | 2.5 | a |
| Short season | V. early | 3.7 | 2.5 | a |
|  | Early | 3.7 | 2.4 | a |
|  | Normal | 3.5 | 1.9 | a |
| Source of variation | df | P>F |  |  |
| Planting date (PD) | 2 | 0.005 | ** |  |
| Hybrid (V) | 1 | <0.001 | *** |  |
| Precipitation levels (dnw) | 2 | <0.001 | *** |  |
| PD x V | 2 | 0.08 |  |  |
| PD x dnw | 4 | <0.001 | *** |  |

**Table S6.** Simulated grain yield (Mg ha^−1^) in different planting dates for a full and short season hybrid in four contrasting sites in Kansas (1986-2015). Each value represents the great mean over four locations and 30 years (1986-2015)

| Location | Planting date | Full season |  | Short season |  |
| --- | --- | --- | --- | --- | --- |
|  |  | mean | sd | mean | sd |
| All | V. early | 4.4 | 2.9 | 3.7 | 2.5 |
|  | Early | 4.4 | 2.9 | 3.7 | 2.3 |
|  | Normal | 3.9 | 2.5 | 3.5 | 1.9 |
| Colby | V. early | 3.7 | 2.8 | 3.0 | 2.7 |
|  | Early | 3.6 | 2.9 | 3.3 | 2.5 |
|  | Normal | 2.6 | 1.8 | 2.9 | 2.0 |
| Garden City | V. early | 2.8 | 2.4 | 2.5 | 2.3 |
|  | Early | 2.9 | 2.3 | 2.9 | 2.4 |
|  | Normal | 2.5 | 1.9 | 2.5 | 1.9 |
| Hayes | V. early | 4.8 | 2.7 | 4.0 | 2.3 |
|  | Early | 5.1 | 2.8 | 4.2 | 2.3 |
|  | Normal | 4.9 | 2.2 | 4.3 | 1.9 |
| Manhattan | V. early | 6.2 | 2.6 | 5.0 | 2.5 |
|  | Early | 6.0 | 2.6 | 4.4 | 1.9 |
|  | Normal | 5.9 | 2.3 | 4.2 | 1.5 |

**Figure S1.**
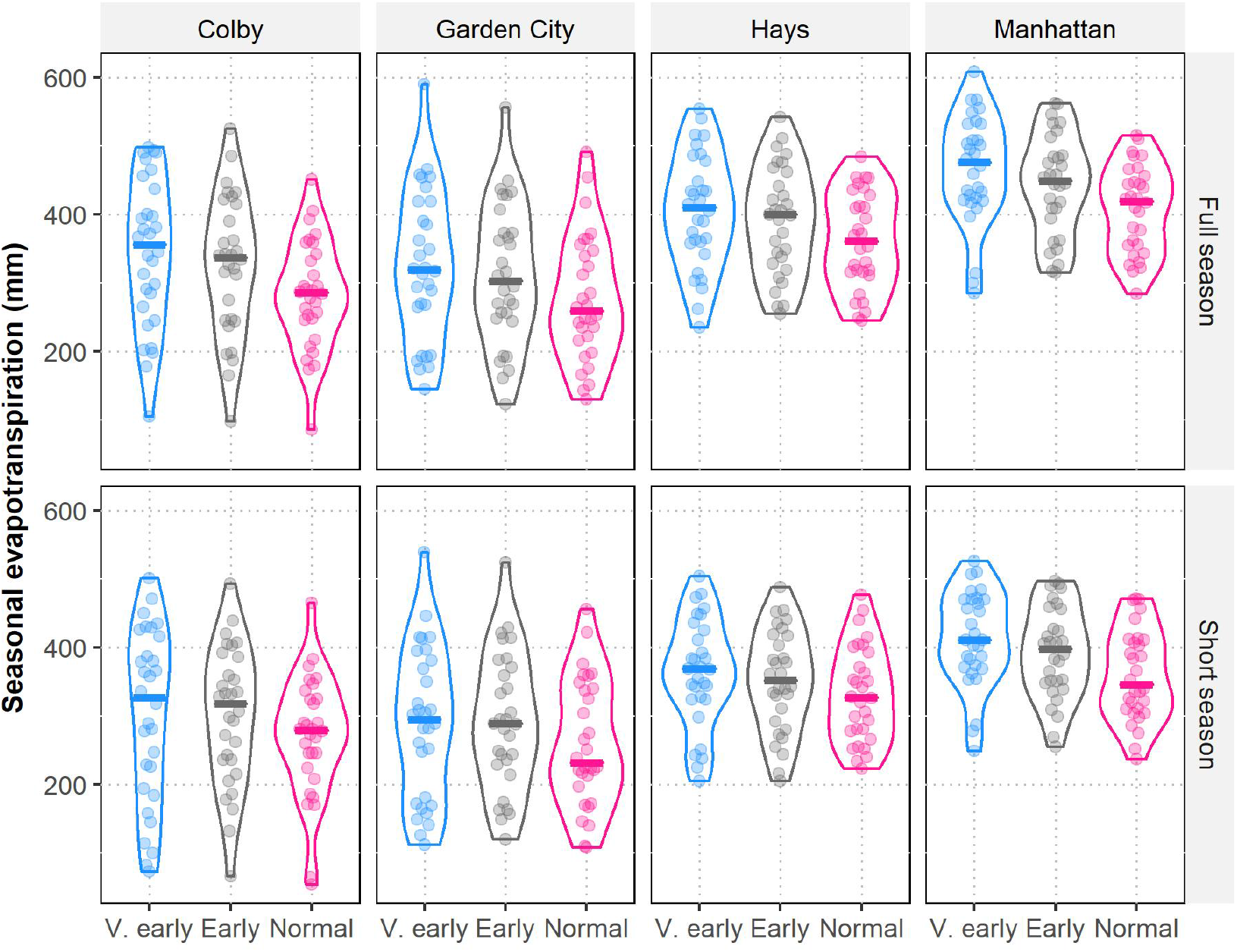
Seasonal evapotranspiration in four contrasting locations and different planting dates for a simulated full-and short-season hybrids. Each violin plot represents the annual variability of 30 years (1986-2015) in Kansas.

**Figure S2.**
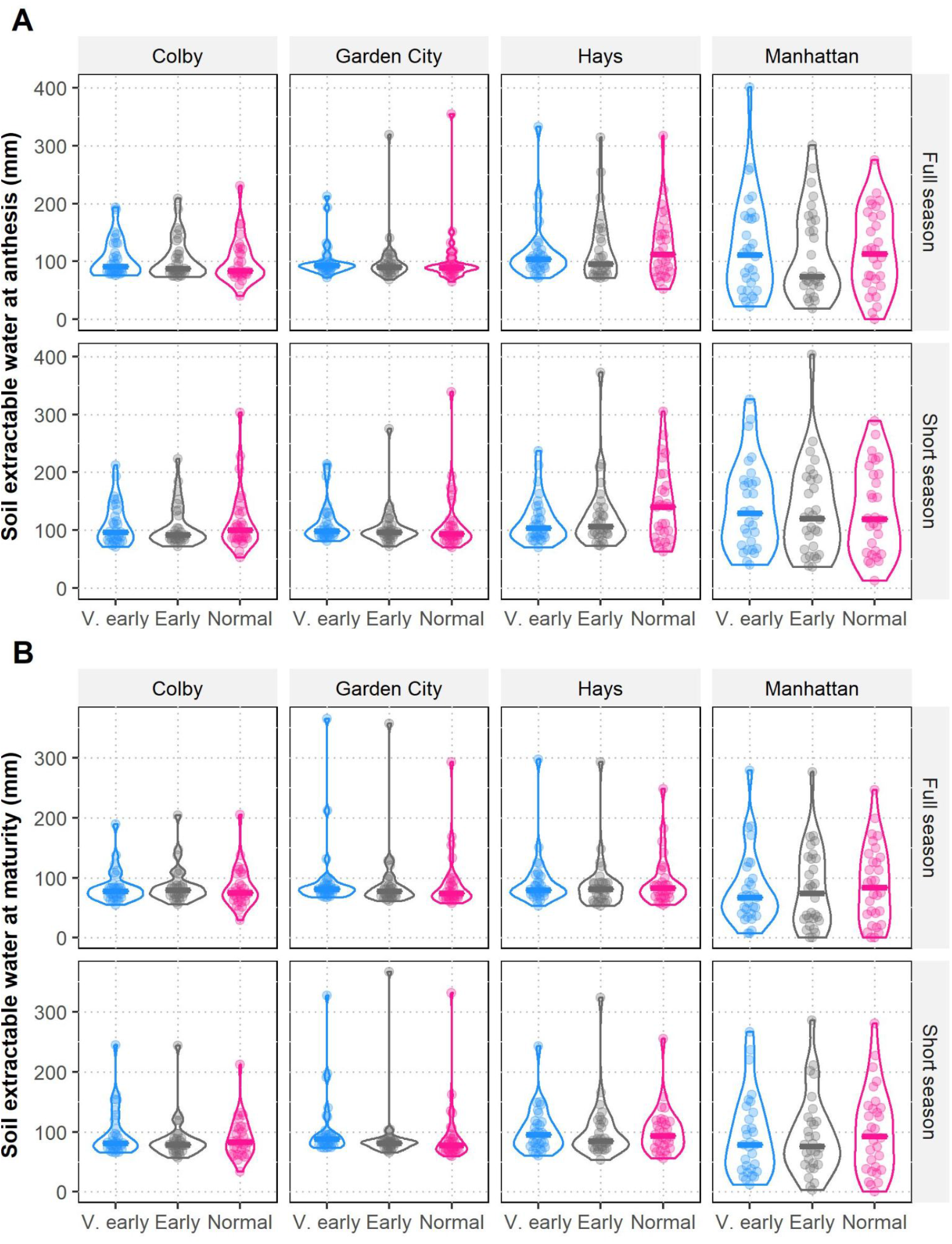
Soil extractable water at (A) anthesis and (B) maturity in four contrasting locations and different planting dates in Kansas for a simulated full-and short-season hybrids. Each violin plot represents the annual variability of 30 years (1986-2015).

## Notes

### Competing Interest Statement

GPM has filed a provisional patent on chilling tolerance markers

### Summary of Updates

- Add more context for current planting dates of sorghum and corn in central US - Add more detailed analysis of effects of base temperate (Tbase)

