## Supplementary File 1 for "Crop modeling defines opportunities and challenges for drought escape, water capture, and yield increase using chilling-tolerant sorghum": Supplementary file 1.docx

### **Script Figure 2**

```
#Figure 2a
rm(list=ls(all=TRUE))
setwd('D:/2019/model_work/calibration')
x<- read.csv("./model_testing_EDAT_chilling tolerance_new.csv",header=TRUE)

ymin<-0
ymax<-20000
xmin<-50
xmax<-150
cex_a<-1
cex_axis<-1.1
tck_<-0.02
cex_m<-0.8
cex_legend<-0.9
cex_mtext <- 1
cex_pch <- 1.8
letter <- 1.5

#dir<-"./model_testing_EDAT_chilling tolerances_new1.jpg"
#jpeg(dir,res=300, width=8.1, height=3, unit="in")

par (mfrow=c(1,3),mar=c(0,0,0,1.5),oma=c(3.5,3.5,2,1), mgp = c(0,0.4,0))

##tbase06
plot(x$O_EDAT,x$S_EDAT_tbase6,pch=(x$number), #main = "1. Tbase: 8 C",
 	ylim=c(0,22),xlim=c(0,22),
 	xlab="", ylab="",axes=FALSE, frame.plot=TRUE, cex=cex_pch)


axis(1, tck=tck_,las=1,cex.axis=cex_axis)
axis(2, tck=tck_,las=1,labels=TRUE,cex.axis=cex_axis)
axis(4, tck=tck_,las=1,labels=FALSE,cex.axis=cex_axis)

segments(-1,-1,30,30)
text(21,22.3,"1:1")
text(15,5,"Tbase: 6 C", cex=1.5, font = 4)
text(15,3,"RMSE = 2.2", cex=1.1)

legend("topleft", xpd = TRUE,legend=x$experiment1,
   	pch =c(1,2,3,4,5,6,7,8), #x.intersp=0.2,
   	horiz=FALSE,
   	cex=cex_legend, bty="n")


##tbase08
plot(x$O_EDAT,x$S_EDAT,pch=(x$number), #main = "1. Tbase: 8 C",
 	ylim=c(0,22),xlim=c(0,22),
 	xlab="", ylab="",axes=FALSE, frame.plot=TRUE, cex=cex_pch)


axis(1, tck=tck_,las=1,cex.axis=cex_axis)
axis(2, tck=tck_,las=1,labels=TRUE,cex.axis=cex_axis)
axis(4, tck=tck_,las=1,labels=FALSE,cex.axis=cex_axis)

segments(-1,-1,30,30)
text(21,22.3,"1:1")
text(15,5,"Tbase: 8 C", cex=1.5, font = 4)
text(15,3,"RMSE = 1.7", cex=1.1)

legend("topleft", xpd = TRUE,legend=x$experiment1,
   	pch =c(1,2,3,4,5,6,7,8), #x.intersp=0.2,
   	horiz=FALSE,
   	cex=cex_legend, bty="n")

##tbase10
plot(x$O_EDAT,x$S_EDAT_tbase10,pch=(x$number), #main = "2. Tbase: 10 C",
 	ylim=c(0,22),xlim=c(0,22),
 	xlab="", ylab="",axes=FALSE, frame.plot=TRUE, cex=cex_pch)


axis(1, tck=tck_,las=1,cex.axis=cex_axis)
axis(2, tck=tck_,las=1,labels=TRUE,cex.axis=cex_axis)
axis(4, tck=tck_,las=1,labels=FALSE,cex.axis=cex_axis)

mtext(text="Observed days to emergence", side = 1, line = 2, outer = TRUE, at = NA,
  	adj = NA, padj = NA, cex =cex_mtext, col = NA)

mtext(text="Simulated days to emergence", side = 2, line = 2, outer = TRUE, at = NA,
  	adj = NA, padj = NA, cex =cex_mtext, col = NA)

mtext(text="a", side = 3,line = 0.3, outer = TRUE, at = 0.015,
  	adj = NA, padj = NA, cex =letter, col = NA)

segments(-1,-1,30,30)
text(21,22.3,"1:1")
text(15,5,"Tbase: 10 C", cex=1.5, font = 4)
text(15,3,"RMSE = 2.0", cex=1.1)


legend("topleft", xpd = TRUE,legend=x$experiment1,
   	pch =c(1,2,3,4,5,6,7,8), #x.intersp=0.2,
   	horiz=FALSE,
   	cex=cex_legend, bty="n")

#dev.off()

#Figure 1b
#remove previous elements
rm(list=ls(all=TRUE))

#set working directory
setwd('D:/tbase_sensibility/')

#main dataframe
summary <-""

#read filed inf the folder
file.names <- dir(pattern =".txt")

file1 <- read.csv("./header_osu.csv", header=TRUE,quote="")

#loop over files, add name to each file and merge to the main dataframe
for(i in 1:length(file.names)){
  name_ <- substring(file.names[i],10,11 )
  file <- read.table(file.names[i],skip=4,header=FALSE,sep="",quote="")
  #read column name of file1
  nam <- names(file1)
  #add column name to file
  colnames(file) <- nam
  file$tbase <-name_
  summary <- rbind(summary, file)
}

#summa <- na.omit(data.frame(summa)) #not
summary <- data.frame(summary[-c(1), ] )
head(summary)
tail(summary)

str(summary)
summary$year <- as.numeric(substring(summary$SDAT,1,4))
summary$loc <- as.factor(substring(summary$WSTA,3,4))
summary$pd <- as.factor(substring(summary$TNAM,10,11))
summary$var <- as.factor(substring(summary$TNAM,7,8))
summary$HWAM <- as.numeric(summary$HWAM)
summary$tbase <- as.numeric(summary$tbase)
summary$planting <- as.numeric(substring(summary$PDAT,5,7))
summary$emergence <- as.numeric(substring(summary$EDAT,5,7))
summary$days_emergence <-summary$emergence-summary$planting

str(summary)

library(tidyverse)
library(dplyr)

#daily summary
summary_mean <- summary %>%
  group_by (loc,pd,var,tbase) %>% # select variables to summarise
  summarise(HWAM1 = mean(HWAM),
        	emergence = mean(days_emergence),
        	emergence_se = sd(days_emergence)/sqrt(30),
        	HWAM_sd = sd(HWAM)
  )

head(summary_mean)
summary_mean$eme_plus<-summary_mean$emergence +summary_mean$emergence_se
summary_mean$eme_minus<-summary_mean$emergence -summary_mean$emergence_se
head(summary_mean)

#add colors to planting dates
summary_mean$col <- ifelse(summary_mean$pd=="p1","black", ifelse(summary_mean$pd=="p5","gray40","gray80"))
#str(summary_mean)

#write.csv(summary_mean, "summary_mean.csv")
loc <- unique(summary_mean$loc)
loc_name <-c("Northwest: Colby","Southwest: G. City","Central: Hays","Northeast: Manhattan")

summary_mean2 <- summary_mean[summary_mean$loc ==loc[1] & summary_mean$var == "11"&summary_mean$pd == "p1",]
model<-lm(summary_mean2$emergence~summary_mean2$tbase)
summary(model)

#dir<-"./tbase_snesibility.jpg"
#jpeg(dir,res=300, width=7.5, height=3, unit="in")

par (mfrow=c(1,4),mar=c(0,0,0,0),oma=c(3.5,3.7,2,0.2), mgp = c(0,0.4,0))
cex_mtext =0.9
cex_mtextT =0.9
letter <- 1.3
cex_text =1.3
cex_legend=1.2
cex =1.1

for(m in 1:4)
{
  panelN <- c("c1.","c2.","c3.","c4.")
  x <- summary_mean[summary_mean$loc ==loc[m] &summary_mean$var == "11",]

  plot(x$tbase,x$emergence, pch=19, col=x$col, cex=1.5,
   	xlab="", ylab="",axes=F,ylim=c(4,20),xlim=c(5.5,10.5))

  # Vertical arrow
  arrows(x0=x$tbase, y0=x$eme_minus, x1=x$tbase, y1=x$eme_plus,
     	code=3, angle=90, length=0.05,
     	col=x$col, lwd=1)
  box()

  #text(6.1,19,panelN[m], cex=cex_text)
  text(5.6,19.8,loc_name[m], cex=cex_text,pos=4,font = 4)

  if(m == 1)
  {
	axis(2, tck=0.02,labels=TRUE,las=1,cex.axis=cex)
	axis(1, tck=0.02,labels=TRUE,las=1,cex.axis=cex)
	axis(4, tck=0.02,labels=FALSE,las=2,cex.axis=cex)
  }
  else
  {
	axis(2, tck=0.02,labels=FALSE,las=1,cex.axis=cex)
	axis(1, tck=0.02,labels=TRUE,las=1,cex.axis=cex)
	axis(4, tck=0.02,labels=FALSE,las=2,cex.axis=cex)
  }
}

mtext(text="Tbase (C)", side = 1, line = 2, outer = TRUE, at = NA,
  	adj = NA, padj = NA, cex =cex_mtextT, col = NA)
mtext(text="Simulated", side = 2, line = 2.7, outer = TRUE, at = NA,
  	adj = NA, padj = NA, cex =cex_mtextT, col = NA)
mtext(text="days to emergence", side = 2, line = 1.65, outer = TRUE, at = NA,
  	adj = NA, padj = NA, cex =cex_mtextT, col = NA)
mtext(text="b", side = 3,line = 0.3, outer = TRUE, at = 0.01,
   	adj = NA, padj = NA, cex =letter, col = NA)
```

#dev.off()

### **Script Figure 3**

```
rm(list=ls(all=TRUE))
setwd('/late_medium_early')

#simulations ID
id <-read.csv("./KGFR8617_ID.csv",header=TRUE)

#Daily soil moisture file
sw <-read.csv("./KGFR8617_OSW1.csv",header=TRUE)

#Substring planting date (PDAT)
DOY <- as.numeric(substr(id$PDAT, 5, 7))
head(DOY)

#merge files
id_ <- data.frame(cbind(id,DOY))
head (id_)

i_sw0 <- merge(id_, sw, by = c("RUNNO","TRNO", "DOY"))
head(i_sw0)

#file with precipitation levels
dnw_year<-read.csv("D:/2019/KS_list_weather_stations/weather/mesonet/precipitation_quartiles/dnw_all.csv",header=TRUE)

#merge with initial soil moisture
i_sw1 <- merge(i_sw0, dnw_year, by = c("locyear","year","LOC"))

#select a full season hybrid
i_sw <- i_sw1[i_sw1$cult == 11, ]

head(i_sw)

###
#convert strings to factors
i_sw$year <-as.factor(i_sw$year)
i_sw$LOC <-as.factor(i_sw$LOC)
i_sw$dnw1 <-as.factor(i_sw$dnw1)
i_sw$cult <-as.factor(i_sw$cult)

#analysis of variance
library(lme4)
library(Matrix)
library(lmerTest)
library(multcomp)
library(car)

####
#run the model with interaction
#SWXD   	<- soil extractable water of soil moisture
#PD     	<- planting date: p1: 15-April,p5: 15-May,p9: 15:June
#dnw2   	<- precipitation levels
#cultivar   <- 11: full season, 21: short season (not simulated)

model.lm <- lmer(SWXD ~ PD + LOC + PD * LOC + (1|year), data=i_sw,REML=F)

summary(model.lm)
anova(model.lm)

#residual analysis
library(ggResidpanel)
resid_panel(model.lm, plots = "all")

#mean comparisons
library(emmeans)
( cld_dat = as.data.frame( cld(emmeans(model.lm, ~ PD |LOC),
                           	Letters = letters ) ) )

#1. Violin plots initial soil moisture
library(ggplot2)
library(gridExtra)
loc <- c("Colby"="Northwest: Colby", "Gcity"= "Southwest: Garden City", "Hayes"= "Central: Hays","Manha"="Northeast: Manhattan")

p <- ggplot(i_sw , aes(x=PD, y=SWXD, color=factor(PD))) + geom_violin(trim=TRUE) +
  	theme(legend.position = "none",
        	plot.background = element_rect(fill = "white"),
        	panel.background = element_rect(fill = "white", colour="black"),
        	panel.grid.major = element_line(colour = "gray", linetype = "dotted"),
        	strip.background = element_rect(fill="gray95"))


library(ggbeeswarm)

p1 <- p + scale_colour_manual(values = rep(c("dodgerblue", "dimgray", "deeppink"),4)) +
  	geom_quasirandom(alpha = 0.3, width = 0.15,size=1.7)+
  	geom_crossbar(stat="summary", fun.y=median, fun.ymax=median, fun.ymin=median, fatten=2.5, width=0.4) +
      scale_x_discrete(labels=c("Early", "Normal", "Late")) +
  	facet_wrap(vars(LOC), ncol = 4,labeller = labeller(LOC = loc)) +
  	geom_text(data = cld_dat, aes(y = 300, label = .group), nudge_x=0.25,color ="black",size=4) +
  	ggtitle("") +
  	xlab("") + xlab("Planting date") +
  	ylab("") + ylab("Extract. water at planting (mm)")

p1

###2. Precipitation at each location
c<-read.csv("weather/Thomas_Colby_1986_2015_f.csv",header=TRUE)
g<-read.csv("weather/Finney_GardenCity_1986_2015_f.csv",header=TRUE)
h<-read.csv("weather/Ellis_Hayes_1986_2015_f.csv",header=TRUE)
m<-read.csv("weather/Ryley_Manhattan_1986_2015_f.csv",header=TRUE)

#merging weather files in a single document
loc <-rep("Colby",10957)
cc <- cbind(loc,c)

loc <-rep("Gcity",10957)
gg <- cbind(loc,g)

loc <-rep("Hayes",10957)
hh <- cbind(loc,h)

loc <-rep("Manha",10957)
mm <- cbind(loc,m)

#x <- rbind(hh,gg,mm,cc)
x <- rbind(cc,gg,hh,mm)
head(x)

data <- within(x,  locyear <- paste(loc, year, sep=""))
head(data)

##merging data
###########################################################
dnw_year<-read.csv("D:/2019/KS_list_weather_stations/weather/mesonet/precipitation_quartiles/dnw_all.csv",header=TRUE)

w.data <- merge(data, dnw_year, by = c("locyear","year"))
w.data1 <- w.data[w.data$loc == "Colby", ]
head(w.data1)


###########################################################
#aggregate
rain <-aggregate(RAIN~loc+year+month+dnw2, data=w.data, sum, na.rm=TRUE)
head(rain)
rain1<-data.frame(rain[1],rain[2],rain[3],rain[4],rain[5])
head(rain1)
newdata <- rain1[order(loc),]
head(newdata)
loc_ <- factor(unique(rain1$loc))

library(ggplot2)
library(gridExtra)

loc1 <- c("Colby"="Northwest: Colby", "Gcity"= "Southwest: Garden City", "Hayes"= "Central: Hays","Manha"="Northeast: Manhattan")

q<- ggplot(rain1 , mapping =aes(x=month, y=RAIN,group=as.factor(month)))+ geom_boxplot(lwd=0.25) + #geom_boxplot(notch=FALSE))+

  theme(legend.position = "none",
    	panel.border = element_rect(fill=NA,colour = "black"),
    	plot.background = element_rect(fill = "white"),
    	panel.background = element_rect(colour="black",fill = "white" ),
    	panel.grid.major = element_line(colour = "gray", linetype = "dotted"),
    	strip.background = element_rect(fill="gray95"))

q1 <- q +
  	geom_dotplot(binaxis='y', binwidth=6, stackdir='center', dotsize=0.3) +
  	stat_summary(fun.y = mean, geom = "point", shape =17, size = 1, color = "red") +
      scale_x_continuous(breaks=c(2,4,6,8,10,12)) +
  	facet_wrap(~loc, ncol = 4,labeller = labeller(loc = loc1), scales="fixed", strip.position="top") +
  	ggtitle("") +
  	ylab("Precipitation (mm)") +
  	xlab("Month")

q1


###.3 Daily soil moisture
##########################################################################

#rm(list=ls(all=TRUE))
setwd('/late_medium_early')

#simulations ID
id <-read.csv("./KGFR8617_ID.csv",header=TRUE)

#Daily soil moisture
sw <-read.csv("./KGFR8617_OSW1.csv",header=TRUE)
head(sw)

sw0 <- data.frame(merge(id, sw, by = c("RUNNO",  "TRNO")))
sw1 <- sw0[sw0$cult == 11, ]

head(sw1)

#precipitation levels
dnw_year<-read.csv("D:/2019/KS_list_weather_stations/weather/mesonet/precipitation_quartiles/dnw_all.csv",header=TRUE)
dnw_year1 <- dnw_year[dnw_year$LOC == "Colby", ]

SW <- merge(sw1, dnw_year, by = c("locyear","year","LOC"))
#SW <- merge(sw1, dnw_year1, by = c("locyear","year"))

tail(SW)

library(ggplot2)
library(gridExtra)

planted <- c(p1="Early", p5="Normal", p9="Late")
loc1 <- c("Colby"="Northwest: Colby", "Gcity"= "Southwest: Garden City", "Hayes"= "Central: Hays","Manha"="Northeast: Manhattan")

r <- ggplot(SW , aes(x=DOY, y=SWXD, color =factor(year))) + geom_line(size=0.1) +
   	theme(legend.position = "none",
      	plot.background = element_rect(fill = "white"),
      	panel.background = element_rect(fill = "white", colour="black"),
      	panel.grid.major = element_line(colour = "gray", linetype = "dotted"),
      	strip.background = element_rect(fill="gray95"))#,

r1 <- r +  facet_grid(PD ~ LOC,labeller = labeller(PD = planted,LOC=loc1), scales="fixed") +
  	geom_vline(data=subset(SW, PD=="p1"), aes(xintercept=105), colour="dodgerblue",linetype="dashed", size=0.8) +
  	geom_vline(data=subset(SW, PD=="p5"), aes(xintercept=135), colour="dimgray",linetype="dashed", size=0.8) +
  	geom_vline(data=subset(SW, PD=="p9"), aes(xintercept=166), colour="deeppink",linetype="dashed", size=0.8) +
  	ylab(expression(paste("Soil extractable water (mm)",sep=""))) +
  	xlab("Julian day")
r1


#merging all figures
library(ggpubr)

q2 <- q1 + theme(plot.margin = unit(c(0, 0, 0.5, 0), "lines"))
r2 <- r1 + theme(plot.margin = unit(c(0.5, 0, -0.15, 0), "lines"))
p2 <- p1 + theme(plot.margin = unit(c(-0.15, 0, 0, 0), "lines"))

figure <- ggarrange(q2, r2,p2,
                	labels = c("a", "b","c"),
                	ncol = 1, nrow = 3,heights=c(1.1,1.7,1.1))
figure

ggsave("CT_paper_figure2_210127.jpeg", plot=figure,width = 6.5, height = 9)
```

### **Script Figure 4**

```
rm(list=ls(all=TRUE))
setwd('/late_medium_early')

#I.------start - weather for each planting date-------------------------
#daily weather data for each year and jplanting date
id <-read.csv("./KGFR8617_OPG-OWE.csv",header=TRUE)

#aggregate daily data and create a dataframe
tmax <- aggregate(TMXD~RUNNO+TRNO+LOC+PD+year+cult+locyear+GS_VR, id, mean)
tmin <- aggregate(TMND~RUNNO+TRNO+LOC+PD+year+cult+locyear+GS_VR, id, mean)
srad <- aggregate(SRAD~RUNNO+TRNO+LOC+PD+year+cult+locyear+GS_VR, id, sum)
pred <- aggregate(PRED~RUNNO+TRNO+LOC+PD+year+cult+locyear+GS_VR, id, sum)
head(tmax[9])

#get only weather data
gs_weather_ <- data.frame( tmax[1],tmax[2],tmax[3],tmax[4],
                       	tmax[5],tmax[6], tmax[7] ,tmax[8],
                       	tmax[9],tmin[9], pred[9], srad[9])

gs_weather <- setNames(gs_weather_, c("RUNNO","TRNO","LOC","PD","year",
                                      "cult","locyear","GS_VR",
                                      "TMXD","TMND","PRED","SRAD"))
head(gs_weather)


head(gs_weather)
#select a sull season hybrid (11: full season and 21: short season)
gs_weather1 <- gs_weather[gs_weather$cult == 11, ]
unique(gs_weather$GS_VR)
gs_weather2_ <- gs_weather1[gs_weather1$LOC != "Gcity", ]
gs_weather3_ <- gs_weather2_[gs_weather2_$LOC != "Hayes", ]

#select only information for emergence date
gs_weather4_ <- gs_weather3_[gs_weather3_$GS_VR == 0, ]

head(gs_weather1)
a<-unique(gs_weather4_$LOC)


####1.1--------------------------weather emergence------------------------------
library(ggplot2)
library(gridExtra)
loc <- c("Colby"="Northwest: Colby", "Manha"="Northeast: Manhattan")
cultivar <- c("21"="early","11"="late")
gs_vr <- c('1' = "Vegetative",'2' = "Reproductive" )

x_text_size =8.5
x_lab_text_size = 9

####1.1.1. maximum temperature
p <- ggplot(gs_weather4_ , aes(x=PD, y=TMXD, color=factor(PD))) + geom_violin(trim=TRUE) +
  	theme(legend.position = "none",
        	plot.background = element_rect(fill = "white"),
        	panel.background = element_rect(fill = "white", colour="black"),
        	panel.grid.major.y =  element_line(colour = "gray", linetype = "dotted", size =0.4),
        	panel.grid.major.x =  element_line(colour = "gray", linetype = "dotted", size =0.4),
        	axis.text.x = element_blank(),#, hjust = 1),
        	axis.text.y = element_text(size = x_text_size, angle = 0),
        	axis.title.y = element_text(color = "black", size = x_lab_text_size, face = "bold", vjust=2.5,margin = margin(t = 0, r = 3, b = 0, l = 0)),
        	axis.title.x = element_text(color = "black", size = x_lab_text_size, face = "bold", vjust=2.5),
        	strip.text.x = element_text(size = x_text_size, colour = "black", angle = 0),
        	strip.background = element_rect(fill="gray95"),
        	panel.border = element_rect(colour = "black", fill=NA))

library(ggbeeswarm)

p1 <- p + scale_colour_manual(values = rep(c("dodgerblue", "dimgray", "deeppink"),4)) +
  	geom_quasirandom(alpha = 0.3, width = 0.15,size=1.7)+
  	geom_crossbar(stat="summary", fun.y=median, fun.ymax=median, fun.ymin=median, fatten=2.5, width=0.4) +
      scale_x_discrete(labels=c("Early", "Normal", "Late")) +
  	coord_cartesian(ylim = c(14, 35)) +
  	facet_wrap(vars(LOC), ncol = 2,labeller = labeller(LOC = loc)) +
  	="black",size=3.5, nudge_x = 0.4) +
  	xlab("") + xlab("") +
  	ylab("") + ylab("Maximum temp. (°C)")

p1

####1.1.2. minimum temperature

a <- ggplot(gs_weather4_ , aes(x=PD, y=TMND, color=factor(PD))) + geom_violin(trim=TRUE) +
  	theme(legend.position = "none",
        	plot.background = element_rect(fill = "white"),
        	panel.background = element_rect(fill = "white", colour="black"),
        	panel.grid.major.y =  element_line(colour = "gray", linetype = "dotted", size =0.4),
        	panel.grid.major.x =  element_line(colour = "gray", linetype = "dotted", size =0.4),
        	axis.text.x = element_blank(),#, hjust = 1),
        	axis.text.y = element_text(size = x_text_size, angle = 0),
        	axis.title.y = element_text(color = "black", size = x_lab_text_size, face = "bold", vjust=2.5,margin = margin(t = 0, r = 3, b = 0, l = 0)),
        	axis.title.x = element_text(color = "black", size = x_lab_text_size, face = "bold", vjust=2.5),
        	#strip.text.x = element_text(size = x_text_size, colour = "black", angle = 0),
        	strip.background = element_rect(fill="gray95"),
        	panel.border = element_rect(colour = "black", fill=NA),
        	strip.text.x = element_blank())

library(ggbeeswarm)

a1 <- a + scale_colour_manual(values = rep(c("dodgerblue", "dimgray", "deeppink"),4)) +
  	geom_quasirandom(alpha = 0.3, width = 0.15,size=1.7)+
  	geom_crossbar(stat="summary", fun.y=median, fun.ymax=median, fun.ymin=median, fatten=2.5, width=0.4) +
  	scale_x_discrete(labels=c("Early", "Normal", "Late")) +
  	facet_wrap(vars(LOC), ncol = 2,labeller = labeller(LOC = loc)) +
  	coord_cartesian(ylim = c(0,23)) +
  	ggtitle("") +
  	xlab("") + xlab("") +
  	ylab("") + ylab("Minimum temp. (°C)")

a1

####1.1.3. precipitation

b <- ggplot(gs_weather4_ , aes(x=PD, y=PRED, color=factor(PD))) + geom_violin(trim=TRUE) +
  	theme(legend.position = "none",
        	plot.background = element_rect(fill = "white"),
        	panel.background = element_rect(fill = "white", colour="black"),
        	panel.grid.major.y =  element_line(colour = "gray", linetype = "dotted", size =0.4),
        	panel.grid.major.x =  element_line(colour = "gray", linetype = "dotted", size =0.4),
        	axis.text.x = element_text(size = x_text_size, angle = 0),#, hjust = 1),
        	axis.text.y = element_text(size = x_text_size, angle = 0),
        	axis.title.y = element_text(color = "black", size = x_lab_text_size, face = "bold", vjust=2.5,margin = margin(t = 0, r = -1, b = 0, l = 0)),
        	axis.title.x = element_text(color = "black", size = x_lab_text_size, face = "bold", vjust=2.5),
        	#strip.text.x = element_text(size = x_text_size, colour = "black", angle = 0),
        	strip.background = element_rect(fill="gray95"),
        	panel.border = element_rect(colour = "black", fill=NA),
        	strip.text.x = element_blank())

library(ggbeeswarm)

b1 <- b + scale_colour_manual(values = rep(c("dodgerblue", "dimgray", "deeppink"),4)) +
  	geom_quasirandom(alpha = 0.3, width = 0.15,size=1.7)+
  	geom_crossbar(stat="summary", fun.y=median, fun.ymax=median, fun.ymin=median, fatten=2.5, width=0.4) +
      scale_x_discrete(labels=c("Early", "Normal", "Late")) +
  	#coord_cartesian(xlim = c(0, 500)) +
 	 facet_wrap(vars(LOC), ncol = 2,labeller = labeller(LOC = loc)) +
  	coord_cartesian(ylim = c(0,600)) +
  	ggtitle("") +
  	xlab("") + xlab("") +
  	ylab("") + ylab("Precipitation (mm)")

b1

e_tn_0 <- p1 + theme(plot.margin = unit(c(0, -1.5, -1,1), "lines"))
a_tx_0 <- a1 + theme(plot.margin = unit(c(-1, -1, -1,1), "lines"))
g_pp_0 <- b1 + theme(plot.margin = unit(c(-1, 0, 0,1), "lines"))
#ws_bar1 <- ws_bar + theme(plot.margin = unit(c(-0.5, 0, 0, 0), "lines"))


####1.2---------------------weather anthesis------------------------------
#select only vegetative stage
gs_weather4_ <- gs_weather3_[gs_weather3_$GS_VR == 1, ]

head(gs_weather1)
a<-unique(gs_weather4_$LOC)

####1.2.1. Maximum temperature
library(ggplot2)
library(gridExtra)
loc <- c("Colby"="Northwest: Colby", "Manha"="Northeast: Manhattan")
cultivar <- c("21"="early","11"="late")
gs_vr <- c('1' = "Vegetative",'2' = "Reproductive" )

x_text_size =8.5
x_lab_text_size = 9

p <- ggplot(gs_weather4_ , aes(x=PD, y=TMXD, color=factor(PD))) + geom_violin(trim=TRUE) +
  theme(legend.position = "none",
    	plot.background = element_rect(fill = "white"),
    	panel.background = element_rect(fill = "white", colour="black"),
    	panel.grid.major.y =  element_line(colour = "gray", linetype = "dotted", size =0.4),
    	panel.grid.major.x =  element_line(colour = "gray", linetype = "dotted", size =0.4),
    	axis.text.x = element_blank(),#, hjust = 1),
    	axis.text.y =  element_text(size = x_text_size, angle = 0),
    	axis.title.y = element_text(color = "black", size = x_lab_text_size, face = "bold", vjust=2.5),
    	axis.title.x = element_text(color = "black", size = x_lab_text_size, face = "bold", vjust=2.5),
    	strip.text.x = element_text(size = x_text_size, colour = "black", angle = 0),
    	strip.background = element_rect(fill="gray95"),
    	panel.border = element_rect(colour = "black", fill=NA))

library(ggbeeswarm)

p1 <- p + scale_colour_manual(values = rep(c("dodgerblue", "dimgray", "deeppink"),4)) +
  geom_quasirandom(alpha = 0.3, width = 0.15,size=1.7)+
  geom_crossbar(stat="summary", fun.y=median, fun.ymax=median, fun.ymin=median, fatten=2.5, width=0.4) +
  scale_x_discrete(labels=c("Early", "Normal", "Late")) +
  #coord_cartesian(xlim = c(0, 500)) +
  facet_wrap(vars(LOC), ncol = 2,labeller = labeller(LOC = loc)) +
  coord_cartesian(ylim = c(14, 35)) +
  ggtitle("") +
  xlab("") + xlab("") +
  ylab("") + ylab("")

p1

####1.2.2. minimum temperature

a <- ggplot(gs_weather4_ , aes(x=PD, y=TMND, color=factor(PD))) + geom_violin(trim=TRUE) +
  theme(legend.position = "none",
    	plot.background = element_rect(fill = "white"),
    	panel.background = element_rect(fill = "white", colour="black"),
    	panel.grid.major.y =  element_line(colour = "gray", linetype = "dotted", size =0.4),
    	panel.grid.major.x =  element_line(colour = "gray", linetype = "dotted", size =0.4),
    	axis.text.x = element_blank(),#, hjust = 1),
    	axis.text.y = element_text(size = x_text_size, angle = 0),
    	axis.title.y = element_text(color = "black", size = x_lab_text_size, face = "bold", vjust=2.5),
    	axis.title.x = element_text(color = "black", size = x_lab_text_size, face = "bold", vjust=2.5),
    	#strip.text.x = element_text(size = x_text_size, colour = "black", angle = 0),
    	strip.background = element_rect(fill="gray95"),
    	panel.border = element_rect(colour = "black", fill=NA),
    	strip.text.x = element_blank())

library(ggbeeswarm)

a1 <- a + scale_colour_manual(values = rep(c("dodgerblue", "dimgray", "deeppink"),4)) +
  geom_quasirandom(alpha = 0.3, width = 0.15,size=1.7)+
  geom_crossbar(stat="summary", fun.y=median, fun.ymax=median, fun.ymin=median, fatten=2.5, width=0.4) +
  scale_x_discrete(labels=c("Early", "Normal", "Late")) +
  facet_wrap(vars(LOC), ncol = 2,labeller = labeller(LOC = loc)) +
  coord_cartesian(ylim = c(0, 23)) +
  ggtitle("") +
  xlab("") + xlab("") +
  ylab("") + ylab("")

a1

####1.2.3. precipitation

b <- ggplot(gs_weather4_ , aes(x=PD, y=PRED, color=factor(PD))) + geom_violin(trim=TRUE) +
  theme(legend.position = "none",
    	plot.background = element_rect(fill = "white"),
    	panel.background = element_rect(fill = "white", colour="black"),
    	panel.grid.major.y =  element_line(colour = "gray", linetype = "dotted", size =0.4),
    	panel.grid.major.x =  element_line(colour = "gray", linetype = "dotted", size =0.4),
    	axis.text.x = element_text(size = x_text_size, angle = 0),#, hjust = 1),
    	axis.text.y = element_text(size = x_text_size, angle = 0),
    	axis.title.y = element_text(color = "black", size = x_lab_text_size, face = "bold", vjust=2.5),
    	axis.title.x = element_text(color = "black", size = x_lab_text_size, face = "bold", vjust=2.5),
    	#strip.text.x = element_text(size = x_text_size, colour = "black", angle = 0),
    	strip.background = element_rect(fill="gray95"),
    	panel.border = element_rect(colour = "black", fill=NA),
    	strip.text.x = element_blank())

library(ggbeeswarm)

b1 <- b + scale_colour_manual(values = rep(c("dodgerblue", "dimgray", "deeppink"),4)) +
  geom_quasirandom(alpha = 0.3, width = 0.15,size=1.7)+
  geom_crossbar(stat="summary", fun.y=median, fun.ymax=median, fun.ymin=median, fatten=2.5, width=0.4) +
  scale_x_discrete(labels=c("Early", "Normal", "Late")) +
  facet_wrap(vars(LOC), ncol = 2,labeller = labeller(LOC = loc)) +
  coord_cartesian(ylim = c(0,600)) +
  ggtitle("") +
  xlab("") + xlab("") +
  ylab("") + ylab("")

b1

e_tn_1 <- p1 + theme(plot.margin = unit(c(0, -1, -1,0), "lines"))
a_tx_1 <- a1 + theme(plot.margin = unit(c(-1, -1, -1,0), "lines"))
g_pp_1 <- b1 + theme(plot.margin = unit(c(-1, 0, 0,0), "lines"))

####1.3---------------------weather grain filling--------------------------------------
#select only reproductive stage
gs_weather4_ <- gs_weather3_[gs_weather3_$GS_VR == 2, ]
library(ggplot2)
library(gridExtra)
loc <- c("Colby"="Northwest: Colby", "Manha"="Northeast: Manhattan")
cultivar <- c("21"="early","11"="late")
gs_vr <- c('1' = "Vegetative",'2' = "Reproductive" )

x_text_size =8.5
x_lab_text_size = 9

####1.3.1. maximum temperature

#p <- ggplot(gs_weather1_ , aes(x=PRED, y=TMXD, color=factor(PD))) + geom_point() +
p <- ggplot(gs_weather4_ , aes(x=PD, y=TMXD, color=factor(PD))) + geom_violin(trim=TRUE) +
  theme(legend.position = "none",
    	plot.background = element_rect(fill = "white"),
    	panel.background = element_rect(fill = "white", colour="black"),
    	panel.grid.major.y =  element_line(colour = "gray", linetype = "dotted", size =0.4),
    	panel.grid.major.x =  element_line(colour = "gray", linetype = "dotted", size =0.4),
    	axis.text.x = element_blank(),#, hjust = 1),
    	axis.text.y = element_text(size = x_text_size, angle = 0),
    	axis.title.y = element_text(color = "black", size = x_lab_text_size, face = "bold", vjust=2.5),
    	axis.title.x = element_text(color = "black", size = x_lab_text_size, face = "bold", vjust=2.5),
    	strip.text.x = element_text(size = x_text_size, colour = "black", angle = 0),
    	strip.background = element_rect(fill="gray95"),
    	panel.border = element_rect(colour = "black", fill=NA))

library(ggbeeswarm)

p1 <- p + scale_colour_manual(values = rep(c("dodgerblue", "dimgray", "deeppink"),4)) +
  geom_quasirandom(alpha = 0.3, width = 0.15,size=1.7)+
  geom_crossbar(stat="summary", fun.y=median, fun.ymax=median, fun.ymin=median, fatten=2.5, width=0.4) +
  scale_x_discrete(labels=c("Early", "Normal", "Late")) +
  facet_wrap(vars(LOC), ncol = 2,labeller = labeller(LOC = loc)) +
  coord_cartesian(ylim = c(14, 35)) +
  ggtitle("") +
  xlab("") + xlab("") +
  ylab("") + ylab("")

p1

####1.3.2. minimum temperature

a <- ggplot(gs_weather4_ , aes(x=PD, y=TMND, color=factor(PD))) + geom_violin(trim=TRUE) +
  theme(legend.position = "none",
        plot.background = element_rect(fill = "white"),
    	panel.background = element_rect(fill = "white", colour="black"),
    	panel.grid.major.y =  element_line(colour = "gray", linetype = "dotted", size =0.4),
    	panel.grid.major.x =  element_line(colour = "gray", linetype = "dotted", size =0.4),
    	axis.text.x = element_blank(),#, hjust = 1),
    	axis.text.y = element_text(size = x_text_size, angle = 0),
    	axis.title.y = element_text(color = "black", size = x_lab_text_size, face = "bold", vjust=2.5),
    	axis.title.x = element_text(color = "black", size = x_lab_text_size, face = "bold", vjust=2.5),
    	#strip.text.x = element_text(size = x_text_size, colour = "black", angle = 0),
    	strip.background = element_rect(fill="gray95"),
    	panel.border = element_rect(colour = "black", fill=NA),
    	strip.text.x = element_blank())

library(ggbeeswarm)

a1 <- a + scale_colour_manual(values = rep(c("dodgerblue", "dimgray", "deeppink"),4)) +
  geom_quasirandom(alpha = 0.3, width = 0.15,size=1.7)+
  geom_crossbar(stat="summary", fun.y=median, fun.ymax=median, fun.ymin=median, fatten=2.5, width=0.4) +
  scale_x_discrete(labels=c("Early", "Normal", "Late")) +
  facet_wrap(vars(LOC), ncol = 2,labeller = labeller(LOC = loc)) +
  coord_cartesian(ylim = c(0, 23)) +
  ggtitle("") +
  xlab("") + xlab("") +
  ylab("") + ylab("")

a1

####1.3.3. precipitation

b <- ggplot(gs_weather4_ , aes(x=PD, y=PRED, color=factor(PD))) + geom_violin(trim=TRUE) +
  theme(legend.position = "none",
    	plot.background = element_rect(fill = "white"),
    	panel.background = element_rect(fill = "white", colour="black"),
    	panel.grid.major.y =  element_line(colour = "gray", linetype = "dotted", size =0.4),
    	panel.grid.major.x =  element_line(colour = "gray", linetype = "dotted", size =0.4),
    	axis.text.x = element_text(size = x_text_size, angle = 0),#, hjust = 1),
    	axis.text.y = element_text(size = x_text_size, angle = 0),
    	axis.title.y = element_text(color = "black", size = x_lab_text_size, face = "bold", vjust=2.5),
    	axis.title.x = element_text(color = "black", size = x_lab_text_size, face = "bold", vjust=2.5),
    	#strip.text.x = element_text(size = x_text_size, colour = "black", angle = 0),
    	strip.background = element_rect(fill="gray95"),
    	panel.border = element_rect(colour = "black", fill=NA),
    	strip.text.x = element_blank())

library(ggbeeswarm)

b1 <- b + scale_colour_manual(values = rep(c("dodgerblue", "dimgray", "deeppink"),4)) +
  geom_quasirandom(alpha = 0.3, width = 0.15,size=1.7)+
  geom_crossbar(stat="summary", fun.y=median, fun.ymax=median, fun.ymin=median, fatten=2.5, width=0.4) +
  scale_x_discrete(labels=c("Early", "Normal", "Late")) +
  #coord_cartesian(xlim = c(0, 500)) +
  facet_wrap(vars(LOC), ncol = 2,labeller = labeller(LOC = loc)) +
  coord_cartesian(ylim = c(0,600)) +
  ggtitle("") +
  xlab("") + xlab("") +
  ylab("") + ylab("")

b1

e_tn_2 <- p1 + theme(plot.margin = unit(c(0, -1, -1,0), "lines"))
a_tx_2 <- a1 + theme(plot.margin = unit(c(-1, -1, -1,0), "lines"))
g_pp_2 <- b1 + theme(plot.margin = unit(c(-1, 0, 0,0), "lines"))
#ws_bar1 <- ws_bar + theme(plot.margin = unit(c(-0.5, 0, 0, 0), "lines"))

library(ggpubr)
library(egg)

#------end - weather for each planting date--------------------------------------

#II.----start - phenology for each planting date----------------------
#summary file
sum__ <-read.csv("./KGFR8617.csv",header=TRUE)
#EDAT: days to emergence

sum00 <- sum__[sum__$LOC != "Gcity", ]
sum_ <- sum00[sum00$LOC != "Hayes", ]

sum0 <- sum_[sum_$cult == 11, ]
head(sum0)

x_text_size =8.5
x_lab_text_size = 9


#1.1. emergence
library(ggplot2)
library(gridExtra)
library(ggbeeswarm)
planted <- c(p1="Early planted", p5="Normal planted", p9="Late planted")
pyears <- c( '1' = "Dry years",'2' = "Normal years",'3' = "Wet years")
loc <- c("Colby"="Northwest: Colby", "Manha"="Northeast: Manhattan")
phenology <- c("anthesis"="Anthesis", "emergence"= "Emergence", "grain filling"= "Grain filling")

p <- ggplot(sum0 , aes(x=PD, y=EDAT1, color=factor(PD))) + geom_violin(trim=TRUE) +
    	ylim(0, 30) +
    	theme(legend.position = "none",

          	plot.background = element_rect(fill = "white"),
          	panel.background = element_rect(fill = "white", colour="black"),
          	panel.grid.major.y =  element_line(colour = "gray", linetype = "dotted", size =0.4),
          	panel.grid.major.x =  element_line(colour = "gray", linetype = "dotted", size =0.4),
          	axis.text.x = element_text(size = x_text_size, angle = 0),
          	axis.text.y = element_text(size = x_text_size, angle = 0),
          	axis.title.y = element_text(color = "black", size = x_lab_text_size, face = "bold", vjust=2.5,margin = margin(t = 0, r = 3, b = 0, l = 0)),
          	axis.title.x = element_text(color = "black", size = x_lab_text_size, face = "bold", vjust=2.5),
          	strip.text.x = element_text(size = x_text_size, colour = "black", angle = 0),
      	    strip.background = element_rect(fill="gray95"),
          	panel.border = element_rect(colour = "black", fill=NA))

p1 <- p + scale_colour_manual(values = rep(c("dodgerblue", "dimgray", "deeppink"),4)) +
  	geom_quasirandom(alpha = 0.3, width = 0.15,size=1.7)+
  	geom_crossbar(stat="summary", fun.y=median, fun.ymax=median, fun.ymin=median, fatten=2.5, width=0.4) +
      scale_x_discrete(labels=c("Early", "Normal", "Late")) +
  	facet_wrap(vars(LOC), ncol = 2,labeller = labeller(LOC = loc)) +
  	ggtitle("Emergence") +
  	theme(plot.title = element_text(hjust = 0.5, face="bold"))+
  	xlab("") + xlab("") +
  	ylab("") + ylab("Number of days")

p1

#1.2. anthesis
z <- ggplot(sum0 , aes(x=PD, y=ADAT1, color=factor(PD))) + geom_violin(trim=TRUE) +
    	theme(legend.position = "none",
          	plot.background = element_rect(fill = "white"),
          	panel.background = element_rect(fill = "white", colour="black"),
          	panel.grid.major.y =  element_line(colour = "gray", linetype = "dotted", size =0.4),
          	panel.grid.major.x =  element_line(colour = "gray", linetype = "dotted", size =0.4),
          	axis.text.x = element_text(size = x_text_size, angle = 0),
          	axis.text.y = element_text(size = x_text_size, angle = 0),
       	   axis.title.y = element_text(color = "black", size = x_lab_text_size, face = "bold", vjust=2.5,margin = margin(t = 0, r = 1, b = 0, l = 0)),
          	axis.title.x = element_text(color = "black", size = x_lab_text_size, face = "bold", vjust=2.5),
              strip.text.x = element_text(size = x_text_size, colour = "black", angle = 0),
          	strip.background = element_rect(fill="gray95"),
          	panel.border = element_rect(colour = "black", fill=NA))

z1 <- z + scale_colour_manual(values = rep(c("dodgerblue", "dimgray", "deeppink"),4)) +
  	geom_quasirandom(alpha = 0.3, width = 0.15,size=1.7) +
  	geom_crossbar(stat="summary", fun.y=median, fun.ymax=median, fun.ymin=median, fatten=2.5, width=0.4) +
  	scale_x_discrete(labels=c("Early", "Normal", "Late")) +
  	facet_wrap(vars(LOC), ncol = 2,labeller = labeller(LOC = loc)) +
  	ggtitle("Vegetative stage") +
  	theme(plot.title = element_text(hjust = 0.5, face="bold"))+
  	xlab("") + xlab("") +
  	ylab("") + ylab("")

z1

#1.3. grain filling

y <- ggplot(sum0 , aes(x=PD, y=GF1, color=factor(PD))) + geom_violin(trim=TRUE) +
    	theme(legend.position = "none",
          	plot.background = element_rect(fill = "white"),
          	panel.background = element_rect(fill = "white", colour="black"),
          	panel.grid.major.y =  element_line(colour = "gray", linetype = "dotted", size =0.4),
          	panel.grid.major.x =  element_line(colour = "gray", linetype = "dotted", size =0.4),
          	axis.text.x = element_text(size = x_text_size, angle = 0),
          	axis.text.y = element_text(size = x_text_size, angle = 0),
          	axis.title.y = element_text(color = "black", size = x_lab_text_size, face = "bold", vjust=2.5,margin = margin(t = 0, r = 1, b = 0, l = 0)),
          	axis.title.x = element_text(color = "black", size = x_lab_text_size, face = "bold", vjust=2.5),
          	strip.text.x = element_text(size = x_text_size, colour = "black", angle = 0),
          	strip.background = element_rect(fill="gray95"),
  	        panel.border = element_rect(colour = "black", fill=NA))

y1 <- y + scale_colour_manual(values = rep(c("dodgerblue", "dimgray", "deeppink"),4)) +
  	geom_quasirandom(alpha = 0.3, width = 0.15,size=1.7) +
  	geom_crossbar(stat="summary", fun.y=median, fun.ymax=median, fun.ymin=median, fatten=2.5, width=0.4) +
      scale_x_discrete(labels=c("Early", "Normal", "Late")) +
  	#coord_cartesian(xlim = c(100, 1000)) +
  	facet_wrap(vars(LOC), ncol = 2,labeller = labeller( LOC = loc)) +
  	ggtitle("Reproductive stage") +
  	theme(plot.title = element_text(hjust = 0.5, face="bold")) +
  	xlab("") + xlab("") +
  	ylab("") + ylab("")

y1
#----end - phenology for each planting date----------------------

library(ggpubr)

figureX <- ggarrange(p1, z1, y1,
                 	nrow = 1)
figureX
#emergence
e_day <- p1 + theme(plot.margin = unit(c(0, -1, -1,1), "lines"))
#anthesis
a_day <- z1 + theme(plot.margin = unit(c(-1, -1, -1,0), "lines"))
#grain filling
g_day <- y1 + theme(plot.margin = unit(c(-1, 0, -1,0), "lines"))
#margin(t = 0, r = 0, b = 0, l = 0, unit = "pt")

library(gridExtra)
figure_pw_all <- ggarrange(e_day, a_day, g_day,
                       	e_tn_0, e_tn_1, e_tn_2,
                       	a_tx_0, a_tx_1, a_tx_2,
                       	g_pp_0, g_pp_1, g_pp_2,
                       	labels = c("a", "b", "c",
                                      "d", "e", "f",
                                      "", "", "",
                                      "", "", ""),
                       	ncol = 3, nrow = 4)

ggsave("weather_colby_manha4.jpeg",plot=figure_pw_all, width = 8.5, height = 7)
```

### **Script Figure 5**

```
rm(list=ls(all=TRUE))

setwd('/late_medium_early/')
#1. total soil evaporation: sorghum seasons + fallow seasons
#____________________________________________________________________________
#This file has soil evaporation for fallow seasons (ESAA)
#ESAA: soil evaporation for fallow seasons
#Source:
# SNX: fallow season before planting
# SQX: fallow season after planting
esaa_fallow  	<- read.csv('/late_medium_early/ET_SQX/ESAA_SNX_SQX.csv',header=TRUE)

#This file has soil evaporation for growing seasons
#ESCP: soil evaporation for growing periods
esaa_grow_season <- read.csv('/late_medium_early/KGFR8617.csv',header=TRUE)

esaa_fallow1 <- aggregate(ESAA~RUNNO+TRNO+YEAR+TNAM+PD+locyear+FA_GS+dnw2, data=esaa_fallow, sum, na.rm=TRUE)
head(esaa_fallow1)
tail(esaa_fallow1)

# rename YEAR by year
names(esaa_fallow1)[names(esaa_fallow1) == "YEAR"] <- "year"

#merge files fo growing and fallow seasons
esaa_total <- merge(esaa_fallow1, esaa_grow_season, by = c("RUNNO","TRNO","TNAM","PD","year"))
head(esaa_total)

#ESCP + ESAA = total soil evaporation
esaa_total$ESAA_total <- esaa_total$ESCP + esaa_total$ESAA
esaa_total$total <- "Total"
#head(esaa_total)
#head(esaa_total[92])

##-----total soil evaporation (growing season + fallow season)------------------------------
library(ggplot2)
library(gridExtra)

x_text_size =10
x_lab_text_size = 16

dnw <- c( '1' = "Dry years",'2' = "Moderate years",'3' = "Wet years")
loc <- c("Colby"="Colby", "Gcity"= "Garden City", "Hayes"= "Hayes","Manha"="Manhattan")

z <- ggplot(esaa_total , aes(x=PD, y=ESAA_total, color=factor(PD))) + geom_violin(trim=TRUE) +
        	theme(legend.position = "none",
              	plot.background = element_rect(fill = "white"),
              	panel.background = element_rect(fill = "white", colour="black"),
              	panel.grid.major.y =  element_line(colour = "gray", linetype = "dotted", size =0.4),
              	panel.grid.major.x =  element_line(colour = "gray", linetype = "dotted", size =0.4),
              	axis.text.x = element_blank(),#, hjust = 1),
              	axis.text.y = element_text(size = x_text_size, angle = 0),
              	strip.text.x = element_text(size = x_text_size, colour = "black", angle = 0),
              	strip.background = element_rect(fill="gray95")
        	)

library(ggbeeswarm)
ev_vio <- z + scale_colour_manual(values = c("dodgerblue", "dimgray", "deeppink")) +
      	geom_quasirandom(alpha = 0.3, width = 0.15,size=1.7) +
      	geom_crossbar(stat="summary", fun.y=median, fun.ymax=median, fun.ymin=median, fatten=2.5, width=0.4) +
      	scale_x_discrete(labels=c("Early", "Normal", "Late")) +
      	facet_wrap(vars( dnw2), labeller = labeller(dnw2 = dnw)) +
      	 ggtitle("") +
      	xlab("") + xlab("") +
      	ylab("") + ylab("")


ev_vio

#processing soil evaporation for growing seasons and fallow periods
##barplots: subsetting
##subsetting original dataset
##fallow period
e_fraction_fallow <- esaa_total[,c(1,2,3,4,5,6,8,9,16)]
head(e_fraction_fallow)
e_fraction_fallow$water_loss <-"Fallow period"
colnames(e_fraction_fallow)[8] <- "soil_evap"

##growing period
e_fraction_grow.season <- esaa_total[,c(1,2,3,4,5,6,8,92,16)]
e_fraction_grow.season$water_loss <-"Growing period"
colnames(e_fraction_grow.season)[8] <- "soil_evap"

e_fraction_all <- rbind.data.frame(e_fraction_fallow,e_fraction_grow.season)

e_fraction_all$total <- "Total"

#evaporation
e_fraction_all_median <- aggregate(soil_evap~PD+water_loss+dnw2, data=e_fraction_all, median, na.rm=TRUE)
farb <- c("dodgerblue", "dimgray", "deeppink")

bar <- ggplot(e_fraction_all_median , aes(x=PD, y=soil_evap)) +
          	geom_bar(stat="identity",aes(fill = PD,alpha=factor(water_loss))) +
          	scale_alpha_manual(values = c(0.5, 0.8)) +
          	scale_fill_manual(values= farb) +
          	theme(legend.position = "none",
                	plot.background = element_rect(fill = "white"),
                	panel.background = element_rect(fill = "white", colour="black"),
                	panel.grid.major.y =  element_line(colour = "gray", linetype = "dotted", size =0.4),
                	panel.grid.major.x =  element_line(colour = "gray", linetype = "dotted", size =0.4),
                	axis.text.x = element_text(size = x_text_size, angle = 0),#, hjust = 1),
                	axis.text.y = element_text(size = x_text_size, angle = 0),
                	axis.title.y = element_text(color = "black", size = x_lab_text_size, face = "bold", vjust=2.5),
                	strip.text.x = element_blank()
                	)

ev_bar <- bar +
      	scale_x_discrete(labels=c("Early", "Normal", "Late")) +
      	facet_wrap(vars(dnw2),labeller = labeller( dnw2=dnw)) +
      	ggtitle("") +
      	xlab("") + xlab("") +
      	ylab("") + ylab("")
ev_bar


##---------------------end - soil evaporation-----------------------------------------------

#2. total transpiration: growing season, vegetative stage and reproductive stage
##---------------------start - transpiration-------------------------------------------------------------------------------------------------

summary <-read.csv("./KGFR8617.csv",header=TRUE)
et <-read.csv("./KGFR8617_OEB_1.csv",header=TRUE)
head(et)

#epaa : plant transpiration
#esaa : soil evaporation

epaa <- aggregate(EPAA~ID1+RUNNO+TRNO+YEAR+TNAM+cult+PD+locyear+GS_VR, data=et, sum, na.rm=TRUE)
head(epaa)

et <- data.frame( epaa[1],epaa[2],epaa[3],epaa[4],
              	epaa[5],epaa[6], epaa[7] ,epaa[8],
              	epaa[9],epaa[10])
et <- setNames(et, c("ID1", "RUNNO","TRNO","YEAR","TNAM","cult",
                 	"PD","locyear","GS_VR","EPAA"))

tail(et)

##levels of precipitation for all locations
dnw_year<-read.csv("D:/2019/KS_list_weather_stations/weather/mesonet/precipitation_quartiles/dnw_all.csv",header=TRUE)

#merge ET file with precipitation levels file
ET <- merge(et, dnw_year, by = "locyear")

tail(ET)
ET1 <- ET[ET$cult == 11, ]
ET1 <- ET1[ET1$EPAA >= 1, ]

library(ggplot2)
library(gridExtra)
library(ggbeeswarm)
planted <- c(p1="Early", p5="Normal", p9="Late")
dnw <- c( '1' = "Dry years",'2' = "Moderate years",'3' = "Wet years")
loc <- c("Colby"="Colby", "Gcity"= "Garcen City", "Hayes"= "Hayes","Manha"="Manhattan")
gs_vr <- c('1' = "Vegetative stage",'2' = "Reproductive stage" )

x_text_size =10
x_lab_text_size = 12

phenology <- c("anthesis"="Anthesis", "emergence"= "Emergence", "grain filling"= "Grain filling")

##fraction of transpiration classified into moisture levels, growing season and planting date
t_fraction_all_median <- aggregate(EPAA~dnw2+GS_VR+PD, data=ET1, median, na.rm=TRUE)

colors#
farb <- c("dodgerblue", "dimgray", "deeppink")

##barplot

bar_ <- ggplot(t_fraction_all_median , aes(x=PD, y=EPAA)) +
          	geom_bar(stat="identity",aes(fill = PD,alpha=factor(GS_VR))) +
          	scale_alpha_manual(values = c(0.5, 0.8)) +
          	scale_fill_manual(values= farb) +
          	theme(legend.position = "none",
                	# legend.title=element_blank(),
                	# legend.text=element_text(size=8),
                	# legend.key.size = unit(0.4, "cm"),
                	plot.background = element_rect(fill = "white"),
                	panel.background = element_rect(fill = "white", colour="black"),
                	panel.grid.major.y =  element_line(colour = "gray", linetype = "dotted", size =0.4),
                	panel.grid.major.x =  element_line(colour = "gray", linetype = "dotted", size =0.4),
                	axis.text.x = element_text(size = x_text_size, angle = 0),#, hjust = 1),
                	axis.text.y = element_text(size = x_text_size, angle = 0),
                	axis.title.y = element_text(color = "black", size = x_lab_text_size, face = "bold", vjust=2.5),
                	strip.text.x = element_blank()
          	)

bar_

tr_bar <- bar_ +
      	scale_x_discrete(labels=c("Early", "Normal", "Late")) +
      	facet_wrap(vars(dnw2),labeller = labeller( dnw2=dnw)) +
      	ggtitle("") +
      	xlab("") + xlab("") +
      	ylab("") + ylab("") +
      	labs(tag = "a2") +
      	coord_cartesian(ylim = c(0, 380))

tr_bar

##total transpiration

SUMMARY <- merge(summary, dnw_year, by = c("LOC","year"))
SUMMARY1 <- SUMMARY[SUMMARY$cult == 11, ]

##to remove outliers
SUMMARY1 <- SUMMARY1[SUMMARY1$EPCM >= 1, ]

q <- ggplot(SUMMARY1 , aes(x=PD, y=EPCM, color=factor(PD))) + geom_violin(trim=TRUE) +
        	theme(legend.position = "none",
              	plot.background = element_rect(fill = "white"),
              	panel.background = element_rect(fill = "white", colour="black"),
              	panel.grid.major.y =  element_line(colour = "gray", linetype = "dotted", size =0.4),
              	panel.grid.major.x =  element_line(colour = "gray", linetype = "dotted", size =0.4),
              	axis.text.x = element_blank(),#, hjust = 1),
              	axis.text.y = element_text(size = x_text_size, angle = 0),
              	axis.title.y = element_text(color = "black", size = x_lab_text_size, face = "bold", vjust=2.5),
              	strip.text.x = element_text(size = x_text_size, colour = "black", angle = 0),
              	strip.background = element_rect(fill="gray95")
        	)

library(ggbeeswarm)

tr_vio <- q + scale_colour_manual(values = c("dodgerblue", "dimgray", "deeppink")) +
      	geom_quasirandom(alpha = 0.3, width = 0.15,size=1.7)+
      	geom_crossbar(stat="summary", fun.y=median, fun.ymax=median, fun.ymin=median, fatten=2.5, width=0.4) +
      	scale_x_discrete(labels=c("Early", "Normal", "Late")) +
      	facet_grid((~ dnw2), labeller = labeller(dnw2 = dnw)) +
      	ggtitle("") +
      	xlab("") + xlab("") +
      	ylab("") + ylab("") +
      	labs(tag = "A1") +
      	# coord_cartesian(xlim = c(1, 1), ylim = c(350, 1), clip = "off") +
      	theme(plot.tag.position = c(.07, .94))

tr_vio


##---------------------end - transpiratio---------------------------------------------------

#3. Water stress processing : growing period, vegetative stage and reproductive stage
##---------------------water stress---------------------------------------------------------

id <-read.csv("./KGFR8617_ID.csv",header=TRUE)
ws_daily <-read.csv("./KGFR8617_OPG1.csv",header=TRUE)
head(ws_daily)

#estimate the minimum water stress
#water stress
#WSPD -> affect photosynthesis
#WSGD -> affect growth

##WS  -> the minimum of WSPD & WSGD
ws_daily$WS  <- pmin(ws_daily$WSPD, ws_daily$WSGD)
head(ws_daily)

#water stress
#WS_6 -> water stress (WS) higher than 0.6
#WS_7 -> water stress (WS) higher than 0.7
#WS_8 -> water stress (WS) higher than 0.8

#count water stress higher than 0.6, 0.7 a& 0.8
ws_daily$WS_6  <- ifelse(ws_daily$WS >= 0.6, 1,0)
ws_daily$WS_7  <- ifelse(ws_daily$WS >= 0.7, 1,0)
ws_daily$WS_8  <- ifelse(ws_daily$WS >= 0.8, 1,0)
head(ws_daily)

##merging OPG file with ID
ws_daily1 <- merge(ws_daily, id, by = c("RUNNO","TRNO"))
head(ws_daily1)

##extract anthesis date (ADAT) for each simulation
ws_daily1$ADAT_DOY  <- substring(ws_daily1$ADAT,5,7)
head(ws_daily1)

#GS_VR: growing season _ vegetative reproductive
#GS_VR: 1 = vegetative, 2 = reproductive
ws_daily1$GS_VR  <- ifelse(ws_daily1$ADAT_DOY >= ws_daily1$DOY, 1,2)
head(ws_daily1)

#aggregating ws for each phenological stage
ws_vr6 <- aggregate(WS_6~RUNNO+TRNO+YEAR+TNAM+cult+PD+locyear+GS_VR, data=ws_daily1, sum, na.rm=TRUE)
ws_vr7 <- aggregate(WS_7~RUNNO+TRNO+YEAR+TNAM+cult+PD+locyear+GS_VR, data=ws_daily1, sum, na.rm=TRUE)
ws_vr8 <- aggregate(WS_8~RUNNO+TRNO+YEAR+TNAM+cult+PD+locyear+GS_VR, data=ws_daily1, sum, na.rm=TRUE)
head(ws_vr6)

ws_vr <- data.frame( ws_vr6[1],ws_vr6[2],ws_vr6[3],ws_vr6[4],
                 	ws_vr6[5],ws_vr6[6], ws_vr6[7] ,ws_vr6[8],
                 	ws_vr6[9],ws_vr7[9], ws_vr8[9])

ws <- setNames(ws_vr, c("RUNNO","TRNO","YEAR","TNAM",
                    	"cult","PD","locyear","GS_VR",
                    	"WS_6","WS_7","WS_8"))
tail(ws)

##merging with dry, moderate and dry years (dnw2)
#dnw_year<-read.csv("D:/2019/KS_list_weather_stations/weather/mesonet/precipitation_quartiles/dnw_all.csv",header=TRUE)

WS <- merge(ws, dnw_year, by = "locyear")

tail(WS)
WS1 <- WS[WS$cult == 11, ]

ws_fraction_all_median <- aggregate(WS_6~dnw2+GS_VR+PD, data=WS1, mean, na.rm=TRUE)

##_______________________________________________________________________________________
## days under water stress

#library(ggplot2)
#library(gridExtra)
planted <- c(p1="Early planted", p5="Normal planted", p9="Late planted")
dnw <- c( '1' = "Dry years",'2' = "Moderate years",'3' = "Wet years")
loc <- c("Colby"="Colby", "Gcity"= "Garden City", "Hayes"= "Hayes","Manha"="Manhattan")
gs_vr <- c('1' = "Vegetative stage",'2' = "Reproductive stage" )

x_text_size =10
x_lab_text_size = 12

phenology <- c("anthesis"="Anthesis", "emergence"= "Emergence", "grain filling"= "Grain filling")

###
farb <- c("dodgerblue", "dimgray", "deeppink")

##Days under water stress during for each vegetative stage
bar__ <- ggplot(ws_fraction_all_median , aes(x=PD, y=WS_6)) +
      	geom_bar(stat="identity",aes(fill = PD,alpha=factor(GS_VR))) +
      	scale_alpha_manual(values = c(0.5, 0.8)) +
      	scale_fill_manual(values= farb) +
      	theme(legend.position = "none",
            	plot.background = element_rect(fill = "white"),
            	panel.background = element_rect(fill = "white", colour="black"),
            	panel.grid.major.y =  element_line(colour = "gray", linetype = "dotted", size =0.4),
            	panel.grid.major.x =  element_line(colour = "gray", linetype = "dotted", size =0.4),
            	axis.text.x = element_text(size = x_text_size, angle = 0),#, hjust = 1),
            	axis.text.y = element_text(size = x_text_size, angle = 0),
            	axis.title.y = element_text(color = "black", size = x_lab_text_size, face = "bold", vjust=2.5),
            	strip.text.x = element_blank()
            	)

bar__

ws_bar <- bar__ +
      	scale_x_discrete(labels=c("Early", "Normal", "Late")) +
      	facet_wrap(vars(dnw2),labeller = labeller( dnw2=dnw)) +
      	ggtitle("") +
      	xlab("") + xlab("") +
      	ylab("") + ylab("") +
      	labs(tag = "c2") +
      	coord_cartesian(ylim = c(1,52.5)) +
      	theme(plot.tag.position = c(.07, .94))

ws_bar

##_____________________________________________________________________________________________
##2...Days under water stress during the growing season
##
#head(ws)

total_ws_vr6 <- aggregate(WS_6~RUNNO+TRNO+YEAR+TNAM+cult+PD+locyear, data=ws, sum, na.rm=TRUE)
head(total_ws_vr6)

dnw_year<-read.csv("D:/2019/KS_list_weather_stations/weather/mesonet/precipitation_quartiles/dnw_all.csv",header=TRUE)

total_WS <- merge(total_ws_vr6 , dnw_year, by = "locyear")

total_WS <- total_WS[total_WS$cult == 11, ]
head(total_WS)

total_WS$Total <-"Total"
head(total_WS)

q_ <- ggplot(total_WS , aes(x=PD, y=WS_6, color=factor(PD))) + geom_violin(trim=TRUE) +
        	theme(legend.position = "none",
              	plot.background = element_rect(fill = "white"),
              	panel.background = element_rect(fill = "white", colour="black"),
              	panel.grid.major.y =  element_line(colour = "gray", linetype = "dotted", size =0.4),
              	panel.grid.major.x =  element_line(colour = "gray", linetype = "dotted", size =0.4),
              	axis.text.x = element_blank(),#, hjust = 1),
              	axis.text.y = element_text(size = x_text_size, angle = 0),
              	axis.title.y = element_text(color = "black", size = x_lab_text_size, face = "bold", vjust=2.5),
              	strip.text.x = element_text(size = x_text_size, colour = "black", angle = 0),
              	strip.background = element_rect(fill="gray95")
        	)

#library(ggbeeswarm)
ws_vio <- q_ + scale_colour_manual(values = c("dodgerblue", "dimgray", "deeppink")) +
      	geom_quasirandom(alpha = 0.3, width = 0.15,size=1.7)+
      	geom_crossbar(stat="summary", fun.y=median, fun.ymax=median, fun.ymin=median, fatten=2.5, width=0.4) +
                	scale_x_discrete(labels=c("Early", "Normal", "Late")) +
      	facet_grid((~ dnw2), labeller = labeller(dnw2 = dnw)) +
      	ggtitle("") +
      	xlab("") + xlab("") +
      	ylab("") + ylab("") +
      	labs(tag = "c1") +
      	theme(plot.tag.position = c(.07, .94))

ws_vio

##---------------------end - water stress ---------------------------------------------------

library(ggpubr)

#Adjusting margings for each component
ev_vio1 <- ev_vio + theme(plot.margin = unit(c(0, 0, -0.5,0), "lines"))
ev_bar1 <- ev_bar + theme(plot.margin = unit(c(-0.5, 0, 0, 0), "lines"))

tr_vio1 <- tr_vio + theme(plot.margin = unit(c(0, 0, -0.5, 0), "lines"))
tr_bar1 <- tr_bar + theme(plot.margin = unit(c(-0.5, 0, 0,0), "lines"))

ws_vio1 <- ws_vio + theme(plot.margin = unit(c(0, 0, -0.5,0), "lines"))
ws_bar1 <- ws_bar + theme(plot.margin = unit(c(-0.5, 0, 0, 0), "lines"))


evapo <-ggarrange(ev_vio1+rremove("xlab"), ev_bar1+rremove("xlab"),
              	labels = c("a",""),
              	#font.label = list(size = 10),#font.label = list(size = 10),
              	#vjust=-3, #hjust = -0.5,
              	ncol=1, heights=c(1,1))

trans <-ggarrange(tr_vio1+rremove("xlab"), tr_bar1+rremove("xlab"),
              	labels = c("b", ""),
              	#vjust=-3, #hjust = -0.5,
              	ncol=1, heights=c(1,1))

wstre <-ggarrange(ws_vio1+rremove("xlab"), ws_bar1+rremove("xlab"),
               	labels = c("c", ""),
               	#vjust=-3, #hjust = -0.5,
               	ncol=1, heights=c(1,1))

evapo1 <- annotate_figure(evapo
                       	right = text_grob("Soil evaporation (mm)", color = "black",face="bold", rot = 90)
)
test

trans1 <- annotate_figure(trans,
                    	right = text_grob("Transpiration (mm)", color = "black",face="bold", rot = 90)
)
test1

wstre1 <- annotate_figure(wstre,
                     	right = text_grob("Water stress events (days)", color = "black",face="bold", rot = 90)
)

water_budgets <-ggarrange(evapo1, trans1, wstre1,ncol=1)
water_budgets

ggsave("water_components_210115_1.jpeg", plot=zz, width = 5.5, height = 9)
```

### **Script Figure 6**

```
rm(list=ls(all=TRUE))
setwd('/late_medium_early')

#reading simulation file
summ <-read.csv("./KGFR8617.csv",header=TRUE)
head(summ)

#levels of precipitation: d: dry, n: moderate, w: wet
#dnw2 column has the classification
dnw_year<-read.csv("D:/2019/KS_list_weather_stations/weather/mesonet/precipitation_quartiles/dnw_all.csv",header=TRUE)

#merging tables
summ1 <- merge(summ, dnw_year, by = c("LOC","year"))

#convert strings to factors
summ1$year <-as.factor(summ1$year)
summ1$dnw2 <-as.factor(summ1$dnw2)
summ1$cult <-as.factor(summ1$cult)

#loading libraries for anova analysis
library(lme4)
library(Matrix)
library(lmerTest)
library(multcomp)
library(car)

#run the model with interaction
#HWAM   	<- soil evaporation during fallow season
#PD     	<- planting date: p1: 15-April,p5: 15-May,p9: 15:June
#dnw2   	<- precipitation levels
#cultivar   <- 11: full season, 21: short season (not simulated)

#model for total HWAM or grain yield
model.lm <- lmer(HWAM ~ PD + cult + dnw2 + PD*dnw2 + (1|LOC) + (1|year), data=summ1,REML=F)

summary(model.lm)
anova(model.lm)

## deviation of the residuals from a normal distribution
library(ggResidpanel)
resid_panel(model.lm, plots = "all")

#mean comparison
library(emmeans)

( cld_dat = as.data.frame( cld(emmeans(model.lm, ~ PD | dnw2:cult),
                           	Letters = letters ) ) )

#load libraries to plot simulations
library(ggplot2)
library(gridExtra)
library(ggbeeswarm)

planted <- c(p1="Early planted", p5="Normal planted", p9="Late planted")
dnw <- c( '1' = "Dry years",'2' = "Moderate years",'3' = "Wet years")
cult1 <- c( '11' = "Full season",'21' = "Short season")

summ1<-summ1[summ1$HWAM >= 2,]

x_text_size =10
x_lab_text_size = 16

p <- ggplot(summ1 , aes(x=PD, y=HWAM/1000, color=factor(PD))) + geom_violin(trim=TRUE) +
       	theme(legend.position = "none",
            	plot.background = element_rect(fill = "white"),
            	panel.background = element_rect(fill = "white", colour="black"),
            	panel.grid.major.y =  element_line(colour = "gray", linetype = "dotted", size =0.4),
            	panel.grid.major.x =  element_line(colour = "gray", linetype = "dotted", size =0.4),
            	axis.text.x = element_text(size = x_text_size, angle = 0),
            	axis.text.y = element_text(size = x_text_size, angle = 0),
            	strip.text = element_text(size = x_text_size, colour = "black", angle = 0),
            	strip.background = element_rect(fill="gray95")
            	)

p1 <- p + scale_colour_manual(values = rep(c("dodgerblue", "dimgray", "deeppink"),4)) +
    	geom_quasirandom(alpha = 0.3, width = 0.15,size=1.7) +
    	geom_crossbar(stat="summary", fun.y=median, fun.ymax=median, fun.ymin=median, fatten=2.5, width=0.4) +
    	scale_x_discrete(labels=c("Early", "Normal", "Late")) +
    	scale_y_continuous(breaks=c(0,2,4,6,8,10,12,14)) +
    	facet_grid((cult~ dnw2), labeller = labeller(dnw2 = dnw, cult= cult1)) +
    	xlab("") + xlab("") +
    	ylab("") + ylab(expression("Grain yield (Mg "~ha^-1~")"))

p1

#save figure file
ggsave("CT_paper_figure5_hwam_late_early_.jpeg", width = 6, height = 5)
```
