## Supplementary File 2 for "Crop modeling defines opportunities and challenges for drought escape, water capture, and yield increase using chilling-tolerant sorghum": read_all.txt

Simulation files
KGFR8617.csv: summary file for all growing season with seasonal environmental variables, biomass, grain yield.
KGFR8617_ID : simulation ID file
KGFR8617_OSW1.csv: daily soil moisture file (30 years, 4 locations x 3 planting dates)
KGFR8617_OPG-OWE.csv: daily weather data for each growing season(30 years, 4 locations x 3 planting dates)
KGFR8617_OEB_1.csv: daily transpiration (30 years, 4 locations x 3 planting dates)
KGFR8617_OPG1.csv: daily plant growth and water stress (30 years, 4 locations x 3 planting dates)
ESAA_SNX_SQX.csv: total soil evaporation for fallow seasons (30 years, 4 locations x 3 planting dates)

Furhter details of the simulations can be 
Codes for Field and Simulated Data can be obtained in the tab "Codes Viewer" of the DSSAT software

Weather files
Ellis_Hayes_1986_2015_f.csv: daily weather file weather for Hays
Finney_GardenCity_1986_2015_f.csv: daily weather file weather for Garden City
Ryley_Manhattan_1986_2015_f.csv: daily wether file for Manhattan
Thomas_Colby_1986_2015_f.csv: Daily weather file for Colby
dnw_all.csv : Annual precipitation at each location and year and precipitation levels (1: dry, 2: moderate, 3: wet)
